# Tissue-wide profiling of human lungs reveals spatial sequestration of macrophages in tuberculosis

**DOI:** 10.1101/2025.04.11.648467

**Authors:** Wei Xiao, Andrew J. Sawyer, Siwei Mo, Xinyu Bai, Qinzhu Yang, Yi Gao, Timothy Fielder, Yue Zhang, Youchao Dai, Qianting Yang, Yi Cai, Guanggui Ding, Guofang Deng, Liang Fu, Camelia Quek, James Wilmott, Umaimainthan Palendira, Warwick J Britton, Daniel L Barber, Joel D Ernst, Ellis Patrick, Carl G. Feng, Xinchun Chen

## Abstract

The immune response to human tuberculosis (TB), particularly in the context of complex lung pathology, remains incompletely understood. Here, we employed whole-slide spatial proteomics to map immune cell organization in TB-affected human lung tissues. Our analysis revealed pronounced spatial segregation of major immune cell populations in non-necrotizing TB lesions. At the tissue level, macrophages and lymphocytes formed distinct cellular communities associated with specific pathological features. At the lesion level, macrophages and B cells showed an inverse relationship in both abundance and spatial distribution. Proinflammatory T cells preferentially accumulated in macrophage-rich lesions but remained largely separated from macrophages. Interestingly, lesions exhibiting clear segregation between T cells and macrophages were more common in subclinical TB than in active disease. These findings suggest that spatial isolation of macrophages from effector lymphocytes may help temper inflammation and potentially prevent lesion progression to necrosis, while also enabling immune evasion by *Mycobacterium tuberculosis*.

**One Sentence Summary:** Xiao et al. reveal spatial segregation of immune cells in TB-lung tissue and link the microenvironmental dynamics to disease states of tuberculosis.

## INTRODUCTION

Tuberculosis (TB), caused by Mycobacterium tuberculosis (Mtb), remains a significant global health challenge with 10.8 million infections and 1.25 million deaths reported in 2023 (*1*). The rise of multidrug-resistant and extensively drug-resistant Mtb strains has further escalated the burden. A deeper understanding of host immune system function is essential for deciphering TB pathogenesis and developing more effective vaccines and host-directed therapies.

TB is a complex human disease characterized by substantial heterogeneity in various levels. The clinical manifestations of TB are highly diverse, rendering the historical binary classification of active versus latent TB inadequate for fully capturing the disease spectrum. A new TB classification framework has recently been developed. This framework encompasses one non-disease state (Mtb infection), two subclinical states, and two clinical states, which are categorized based on macroscopic pathology, infectiousness, and associated symptoms or signs (*2*). Moreover, the host immune response to Mtb infection exhibits considerable heterogeneity. Mtb is an intracellular bacterium believed to primarily induce a cell-mediated immune response (*3–5*). However, the Mtb-specific antibody response has been linked to enhanced resistance to Mtb infection (*6, 7*) and B cells have been found in the lungs of TB patients (*8–11*). How these functional distinct immune responses coordinated in lung tissue is known.

In addition, there is substantial heterogeneity in the histopathology of human TB lung tissues (*12–18*). Presentations can range from non-necrotizing cell aggregates to more advanced tissue-damaging changes, such as caseous necrosis and cavitation. In this regard, focusing on specific lesion types, such as necrotic or caseating granulomas, risks overlooking the cellular dynamics that lead to terminal tissue pathology, and may be less useful in elucidating the relationship between tissue pathologies and the broad spectrum of TB disease manifestations. To fully capture the complexity of TB pathology, a high-resolution analytical approach that preserves spatial context across the full spectrum of lesion types is essential.

A recent single-cell spatial proteomics study using Multiplexed ion beam imaging (MIBI) with 37-plex antibody panel imaging selected regions of interest has clearly demonstrated the diversity in the immune cell composition of cellular aggregates (*8*). To capture the histopathological terrains across the entire tissue and analyze cellular structures in their entirety, we previously opted to use a tissue-wide mapping approach with 6-plex protein markers and developed a method capable of stratifying TB lesions based on cell composition and spatial organization of immune cells (*9*). While this approach was useful in defining diverse types of TB lesions, it had limited power in deciphering the mechanisms underpinning cell-cell interplay due to the low number of immune lineage markers in the assay.

In this study, we utilized PhenoCycler Fusion (PCF) imaging technology with a comprehensive 35-plex antibody panel, which included a broad selection of immune and non-immune cell markers, to map the immune landscape of entire lung tissue sections from 22 TB patients. This system allowed for the visualization of a wide range of pathological structures, including large TB cavities, at a cellular level. We developed a deep-learning-assisted strategy for lesion identification, coupled with a hierarchical analysis pipeline to investigate cellular responses at both the whole tissue level and within multicellular neighborhoods. Using this approach, we successfully extracted over 525 lesions from 10 whole-slide lung images with minimal selection bias. Our findings demonstrate that distinct immune responses operate concurrently in a highly organized manner across and within TB lesions. We uncovered that macrophages are segregated from both T and B lymphocytes in non-necrotizing TB lesions. Interestingly, compared to clinical TB cases, subclinical patients showed an increased number of lesions where T cells and macrophages were distinctly segregated. Together, these findings uncover an unexpected spatial arrangement of immune cells in TB-affected lungs, potentially serving as a key mechanism for restraining excessive inflammation within diseased tissue.

## RESULTS

### Multiplexed whole-slide imaging captures the landscape of human TB immunopathology with cellular resolution

The histopathological features of TB lung tissues are known to be complex and heterogenous (*8, 9, 12, 14, 16*) (Fig. 1A). To capture and deconvolve the unique pathological landscape with single-cell resolution, we performed whole-slide imaging of twenty-two surgically-resected human lung tissue section using the PCF (Akoya). The 35-plex panel included markers for major cell types such as lymphocytes, myeloid cells, epithelial cells, and endothelial cells, along with 13 other immune markers including PD-1, PD-L1, ICOS and Indoleamine 2,3-dioxygenase 1 (IDO1) (fig. S1A). Following imaging, the same slides were subjected to Hematoxylin and Eosin (H&E) staining (Fig. 1B, fig. S1B). This approach allowed us to link distinct cellular structures defined by the multiplexed antibody staining to those annotated histopathologically based on H&E staining.

**Fig. 1.**
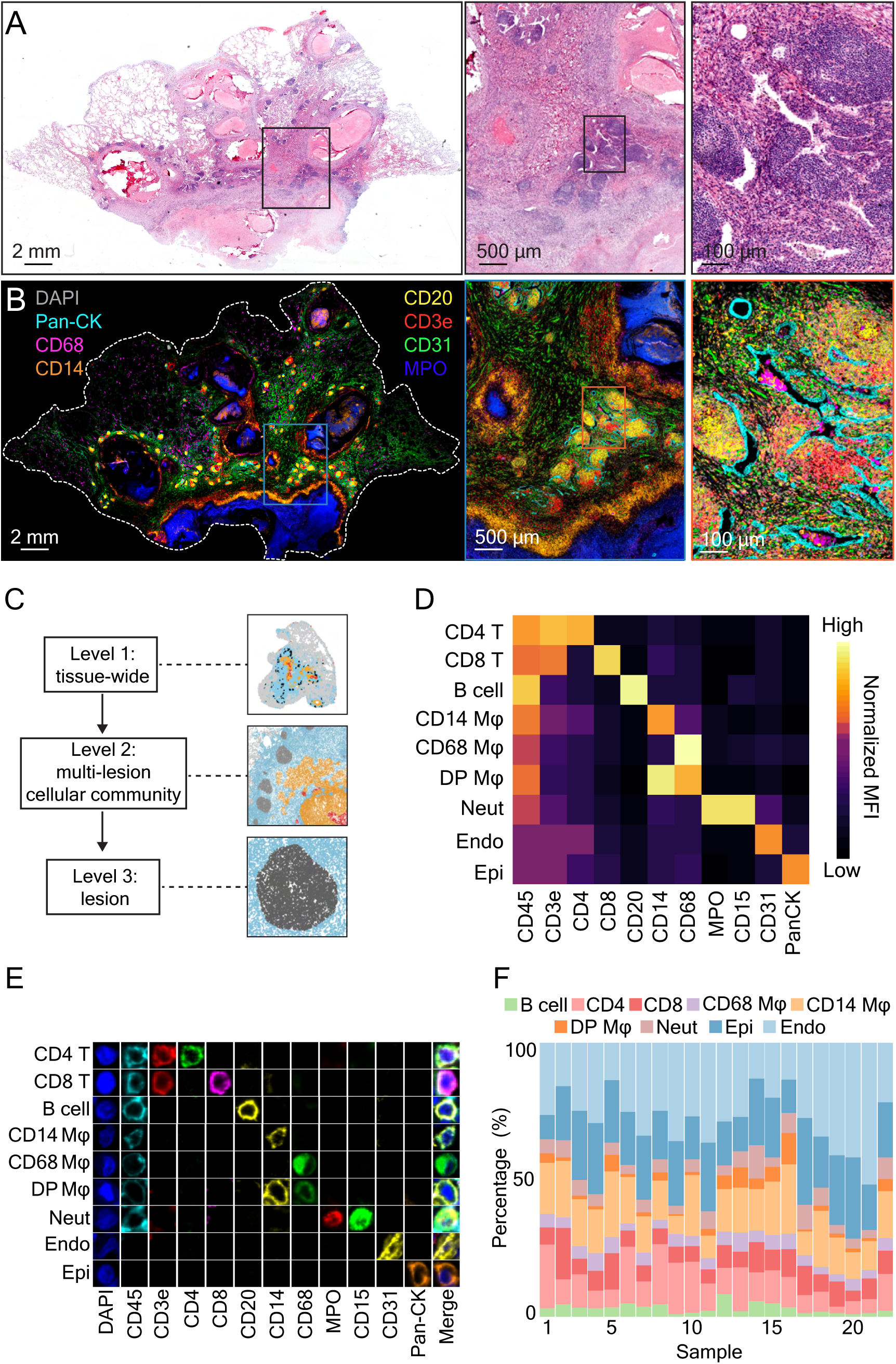
Spatial proteomic profiling of human TB lung tissues using a hierarchical analysis pipeline. Representative **(A)** H&E and **(B)** PCF immunofluorescence images of the same human TB lung section. Boxed regions are magnified in the immediate images on the right. Selected cellular markers are indicated in panel (B). **(C)** Schematic of hierarchy analysis workflow. **(D)** Heatmap showing lineage annotations of the cells from 22 samples based on normalized expression of lineage markers (heatmap columns). **(E)** Imaging validation of lineage identity by overlaying fluorescence channels of DAPI and lineage markers. Rows and columns represent cell types and lineage markers, respectively. **(F)** Percentages of major cell populations among total segmented cells colored by cell type. Each column represents an individual clinical sample.

The 22 specimens represented two clinically confirmed TB patient groups. Clinical TB group comprised patients who had presented with common symptoms of active TB infection (including cough, blood in sputum, fever), tested positive for Mtb pathogen, and undergone anti-TB drug treatment with varying durations prior to lung resection surgery (table S1, patient ID 1-11). Subclinical group consisted of individuals who displayed no typical TB symptoms but showed chest CT abnormality, which were initially suspected to be malignant but subsequently diagnosed as TB during post-surgery histopathological examination (table S1, patient ID 12-22). Mtb bacilli were detected in 3 of 11 lung sections of sub-clinical TB cases. The lung specimens were evaluated independently by two clinical pathologists.

High-resolution imaging of whole tissue sections enabled comprehensive characterization at both the tissue-wide scale and the level of individual lesions with single-cell resolution. We developed a hierarchical analytic strategy for examining multi-lesional cellular community and multi-cellular lesions including necrotic or non-necrotic types (Fig. 1C). The StarDist deep learning segmentation tool (*19*) was employed to identify individual cells within tissues. This segmentation across the 22 tissue samples yielded a total of 20.6 million cells, with each sample containing an average of 934,966 cells (s.d. = 453,773).

Using FuseSOM (*20*), we identified nine major cell populations within the tissue based on the differential expression of leukocyte lineage and stromal cell markers (Fig. 1D). Lymphocytes were identified through the expression of markers CD45, CD3e, CD4, CD8, and CD20. In the myeloid compartment, macrophages were characterized by the expression of CD14 and CD68, while neutrophils were defined by CD15 and MPO expression. Endothelial and epithelial cells were identified using Pan-Cytokeratin (PanCK) and CD31, respectively. The accuracy of cell type annotations was validated by overlaying cell labels onto raw immunofluorescence images alongside the respective marker channels and determining visually whether the staining pattern matched each annotated cell type (Fig. 1E). While varying in their percentages, all cell populations were identified in all 22 samples analyzed in this study (Fig. 1F). The most abundant immune cell populations in the human TB lung sections were CD14^+^ macrophages (mean = 15.90%, s.d. = 5.07% of total annotated cells) and CD4^+^ T cells (11.61% ± 5.94%), followed by CD8^+^ T cells (7.91% ± 3.35%) and B cells (3.14% ± 1.85%). Other myeloid cells included CD68^+^ macrophages (4.41% ± 1.18%), CD14^+^CD68^+^ macrophages (3.02% ± 2.73%), and neutrophils (5.41% ± 2.23%). Non-immune cells were mainly endothelial cells (27.86% ± 9.9%) and epithelial cells (20.75% ± 5.71%), which form the structural framework of lung tissue. Collectively, these findings demonstrate that all major immune cell types are present in human TB lung lesions, with percentages ranging from 0.45% to 51.68% across our dataset.

### Hierarchical stratifying cellular communities and lesions reveals spatial separation of B cells and macrophages

A diverse array of histopathological features identified on whole H&E-stained slides are commonly used for TB diagnosis clinically. To determine tissue-wide TB histopathological features, the same tissue sections used for PCF imaging were H&E-stained and annotated by clinical pathologist. The pathologist-curated annotations were then used to train a deep-learning-based segmentation model (Fig. 2A). This model successfully identified four primary pathological patterns, including histiocytic cuff (HC) (ring-shape aggregation of histiocytes encircling necrotic granulomas), necrosis (NC), lymphoid aggregates (LA), and healthy alveoli (HA) with performance level similar to human pathologist reviews.

**Fig. 2.**
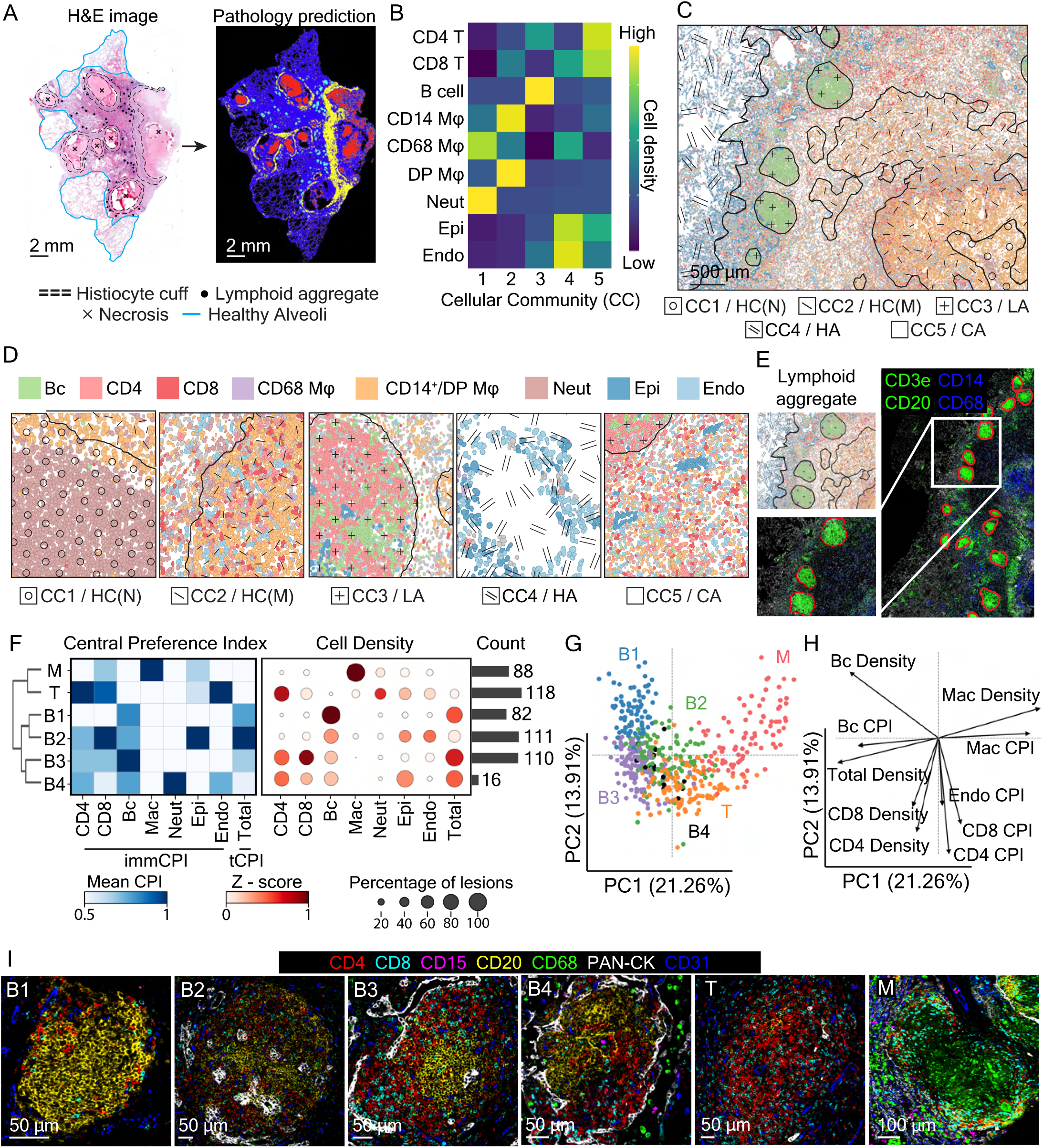
Defining tissue-wide cellular communities and stratifying sub-community non-necrotizing lesions. Cellular-community (CC) defined based on the distribution of the major cell populations **(A)** Schematic workflow for histopathologically defined features on the post-PCF tissue section. **(B)** Heatmap showing the preference of major cell populations (normalized mean density of proportion) in each CC. **(C)** Representative CC phenotype map showing distribution of defined CCs. HC(N), histiocytic cuff (neutrophil enriched); HC(M), histiocytic cuff (macrophage enriched); LA, lymphoid aggregates; HA, healthy alveoli; CA, consolidated area. **(D)** Representative CC phenotype maps showing corresponding field of views of individual CC. **(E)** Schematic diagram showing the identification and extraction of cell aggregate using SAM model with major leukocyte marker and DAPI input. **(F)** Heatmap showing lesion classification using the combination of absolute total and immune cell central preference index (left) and normalized cell type density (right). The horizontal bars indicate the numbers of each lesion type. **(G)** PCA plot visualization of the variations across all aggregates (*n* = 525). Each dot represents an individual cell aggregate and each color corresponds to one defined lesion type. **(H)** PCA loading plot representing the top 10 contributing variables to the PCA plot in (G). **(I)** Representative PCF images of classified lesion types with selected lineage markers (CD4, CD8, CD15, CD20, CD31, CD68, and Pan-CK) shown.

However, H&E-stained images do not provide detailed cellular information. To better characterize the variation in cell type composition of various TB pathologies, we adapted an unsupervised method (*21*) to analyze 22 PCF images. This method involved capturing the abundance of neighboring cell types within a 50 μm radius of each cell and grouping the cells into distinct cellular communities based on their closest neighbors, generating a comprehensive cellular community (CC) map for each TB-affected lung tissue. Five distinct cellular communities were identified within TB lungs, each characterized by unique combinations of cell types and spatial distribution (Fig. 2B). The community masks were mapped back to the immunofluorescence images, allowing for examination of the immune landscape within the tissue context (Fig. 2C). We observed that these communities were present in all 22 tissue sections examined, although the proportions of each community varied among samples (fig. S2A and S2B), indicating that each of these regional cellular responses are common features of human pulmonary TB.

The myeloid-enriched community (CC1), dominated by neutrophils and CD68⁺ macrophages, constituted 1.85% (±1.57%) of total annotated CC at the histiocytic cuff (HC) bordering necrotic lesions, often surrounded by another myeloid-rich community (CC2) at 10.56% (±4.62%), primarily consisting of CD14⁺CD68⁺ macrophages (Fig. 2C and 2D, fig. S2C). A lymphoid aggregate community (CC3), containing both B and T cells, appeared as small clusters throughout the tissue, accounting for an average of 5.48% (±2.55). In well-ventilated areas marked by intact alveolar architecture and minimal inflammation, a stromal cell–driven community (CC4) represented for 47.18% (±9.51) of the annotated CC, featuring CD68⁺ alveolar macrophages, PanCK⁺ epithelial cells, and CD31⁺ endothelial cells. Finally, an additional community (CC5), rich in CD4⁺ and CD8⁺ T cells, represented 34.93% (±8.27) in consolidation areas characterized by diffuse leukocyte infiltration.

To relate pathology patterns to spatial tissue architecture, we compared the pathology model predictions against the CC-map across six samples, yielding a median Pearson R score of 0.303 (fig. S2D and S2E). CC1 and CC2 were found to align with histiocytic cuff, and CC3 matched the scattered lymphoid aggregates. CC4 aligned effectively with the uninfected alveoli regions. Among these, CC3 and CC4 had the strongest correlation with pathologists’ annotations. Of note, CC5 represents an intermediate area between the lesions that were not annotated by the pathologists. These findings provide a cellular basis for TB pathology at the whole tissue level and serve as a guide for downstream capturing non-necrotic lesions for cellular analysis.

To define the boundaries of non-necrotic lesions, we employed a Large Vision Model (LVM) (*22*) enhanced segmentation tool that relies on graph-based input from key leukocyte lineage markers (CD3e, CD20, CD14, CD68) and nuclei marker (DAPI) for cell morphology. Specifically, these marker channels were provided to the model for guiding the automated lesion detection and delineation of individual lesion boundaries (*23*) (Fig. 2E). The identified aggregates were further validated by cross-referencing tissue CC maps and corresponding H&E images. This approach allowed us to identify and extract 525 lesions from 10 tissue samples.

The extracted lesions were then stratified based on the cellular composition and organization of individual lesion as previously described (*9*). Briefly, in addition to assessing the cell density of major cell types within the lesions, we incorporated two spatial evaluation indices into our lesion clustering strategy: total Central Preference Index (tCPI) and Immune Cell Preference Index (immCPI) to measure lesion compactness and cell phenotype centrality (*9*). The tCPI measures the overall compactness of the lesion, while immCPI determines the centrality of each major cell population. Unsupervised clustering of cell type density, tCPI, and immCPI across cell types within each lesion identified six distinct lesion types with diverse cellular composition and organization, suggesting the presence of distinct localized immune responses across TB-affected lung tissues (Fig. 2F).

The use of a high-plex image platform in the current study enabled us to refine to types of non-necrotic lesions, from 4 described previously (*9*) to 6. Among the 525 lesions, 319 (61%) were identified as B cell-driven aggregates with varying composition and spatial organization, these lesions were given a group name beginning with “B” (Fig. 2G and 2H). B1 lesions were primarily composed of highly compacted B cells (Fig. 2I). B2 lesions featured epithelial cells at their core, suggesting a lymphoid enriched peribronchial inflammation. B3 lesions, resembling tertiary lymphoid structures (TLS), contained a B cell center surrounded by both CD4⁺ and CD8⁺ T cells. A minor subset of lesions (B4) was characterized by epithelial cells encasing a T and B lymphocyte mix with small numbers of centralized neutrophils, corresponding to areas of inflammation within intact alveoli. Additionally, we identified two myeloid-dominant lesion types. T type lesions were loosely organized cellular mixtures containing T cells and various other immune cell populations. M type lesions were composed predominantly of macrophages forming a dense myeloid core surrounded by T cells.

Consistent with our previous study (*9*), principal component analysis (PCA) of these lesions revealed that instead of forming distinctive clusters, lesions were found to be organized into a continuous spectrum, with the extremes of the variation driven by B cell and macrophage density. The intermediate lesions displayed heterogeneous spatial patterns (B2-B4 and T types), indicating complex and heterogenous immune cell interactions operating in these types of lesions (Fig. 2G-I). Together, these indicate a spatial separation between B cells and macrophages in non-necrotic TB lesions. However, it should be noted that in this study, lesion grouping is used as a practical tool to support quantitative lesion-level analysis, without implying strict or absolute separation between the groups.

### Functionally diverse T cell subsets are distributed differentially across non-necrotic lesions

To define the organization of T lymphocytes in human TB lungs, we used nine T cell function-related markers to sub-divide CD4^+^ and CD8^+^ T cells into sub-clusters with diverse functional marker expressions. Unsupervised Leiden-clustering (*24*) revealed ten CD4^+^ T cell clusters (C01–C10) and five CD8^+^ T cell clusters (C11–C15), each exhibiting distinct immune phenotypes (Fig. 3A).

**Fig. 3.**
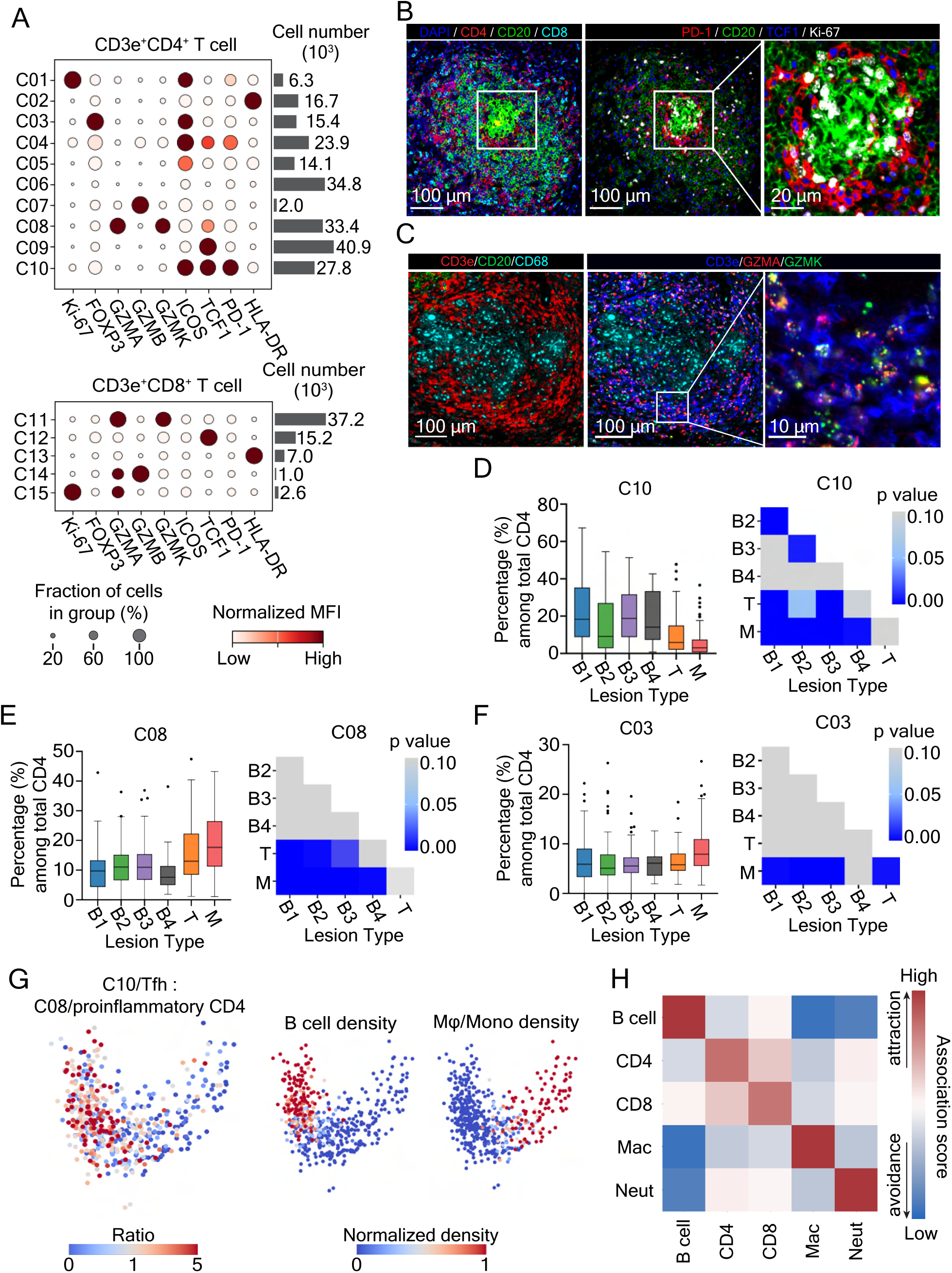
T cell phenotypes and spatial localization are diverse across lesions in TB lung tissue. **(A)** Dotplot graph showing CD4^+^ (top) and CD8^+^ (bottom) T cell sub-populations clustered based on normalized mean fluorescence intensity (MFI) expression of indicated markers. Rows are ordered based on the similarity in marker expression patterns. **(B)** Representative PCF images of Tfh cells in a lymphoid aggregate showing expression of indicated markers. Left image: CD4, CD8, and CD20. Middle and right images: PD-1, TCF1, and Ki-67. The boxed areas in the left and middle images highlight the same region. The right panel displays a magnified view of this specific region. **(C)** Representative PCF images of a myeloid cell-enriched lesion with lymphocytic cuff containing granzyme expressing T cells. Left image: CD3e, CD20, and CD68. Middle and right images: CD3e, GZMA and GZMK. The right panel displays a magnified view of the boxed region in the middle image. **(D -F)** Box plot (left) showing the percentage of **(D)** Tfh (C10), **(E)** Treg (C03) and **(F)** proinflammatory CD4^+^ T cell (C08) among total CD4^+^ T cells per lesion across all lesion types, and heatmap (right) indicating statistical significance from post hoc multi-group comparison for each respective cluster with *P* value shown. The level of statistical significance is determined using One-way ANOVA, the level of significance was set at *P* = 0.05. **(G)** PCA projection of lesion distribution as Fig. 2G colored by raw values of Tfh (C10) to proinflammatory CD4^+^ T cell (C08) ratio in individual aggregate (left), normalized B cell and macrophage density (right). **(H)** Heatmap indicating the median spatial association scores derived from the proximity between major immune populations among all the lesions (*n* = 525).

Among the CD4^+^ T cell clusters, C01 showed high expression level of Ki-67 and ICOS, indicative of proliferating helper T cells. C02 expressed high levels of HLA-DR, putatively identifying as activated, antigen-experienced cells. C03 expressed FOXP3 and ICOS, indicative of regulatory T cells in tissues. C04 and C05 displayed varying levels of ICOS without expression of other T cell markers. C06 did not express any markers examined here, suggesting a CD4⁺ T cell population that requires further characterization. C07 represented a small subset of GZMB-expressing cells while C08 contained a large population of GZMA and GZMK-expressing cells. C09 expressed TCF1, a marker that has been previously proposed to indicate progenitor-like T cells (*25–27*) although TCF1 can potentially be expressed in a subset of Th17 cells (*28*). Additionally, we identified cells in C10 were positive for ICOS, TCF1, and PD-1, suggesting a follicular helper T cell (Tfh) phenotype.

Five CD8^+^ clusters were identified: C11 (GZMA^+^ and GZMK^+^), resembling a recently identified proinflammatory CD8^+^ T cell population (*29, 30*); C12 (TCF1^+^), progenitor-like cells; C13 (HLA-DR^+^), activated CD8⁺ T cells; C14 (GZMA^+^ and GZMB^+^), cytotoxic cells; and C15 (Ki-67^+^), proliferating CD8⁺ T cells.

To explore whether some T cell subsets are preferentially associated with certain lesion types, we focused on Tfh cells (C10) and proinflammatory CD4^+^ T cell subsets (C08, termed based on the recently identified granzyme K function). Cell phenotype map revealed that Tfh cells were enriched in B-type lesions accumulating at the border of germinal center-like structures in B3 type lesions (Fig. 3B). The expression levels of TCF1 and PD-1 varied across 319 B-type lesions were negatively correlated (Spearman *r* = – 0.46, *P* < 1×10⁻⁴) (fig. S3A), revealing that the differentiation state of Tfh in tissue is highly dynamic and lesion-dependent. For proinflammatory cells, co-expressions of GZMA and GZMK were observed in both CD4^+^ and CD8^+^ T cells within the lesions (Fig. 3A and 3C). Importantly, our composition analysis demonstrated that these T cell subsets, representing distinct immune responses, were differentially enriched depending on lesion types. Tfh cells were predominantly found in B-type lesions (Fig. 3D), while proinflammatory CD4^+^ T cells resided mainly in M and T-type lesions (Fig. 3E). Interestingly, regulatory T cells were enriched in the former lesion type (Fig. 3F). Interestingly, there was no lesion-specific enrichment of proinflammatory CD8^+^ T cells across lesion types (fig.S3B). When the 525 lesions were color-coded based on the ratio of Tfh cells to proinflammatory CD4⁺ T cells in each lesion, we observed that along the continuum of lesions, Tfh cells were preferentially enriched in B-type lesions, whereas cytotoxic CD4^+^ T cells were highly enriched in lesions with abundant macrophages (Fig. 3G).

Finally, to evaluate the spatial association between lesional cells along these continuum, we employed a spatial proximity analysis performing inference on cell-type co-localization (*31*). We observed that B cells tend to be located further away from macrophages within the microenvironment, with minimal preference towards or away from other immune cell populations (Fig. 3H), suggesting an inverse relationship in the spatial distribution of B cells and macrophages in the non-necrotizing lesions.

### Immune cell populations are highly organized within non-necrotizing TB lesions

To gain a systematic view of T cell distribution across lesion types, we quantified each T cell clusters in aggregated data for each lesion type (Fig. 4A). This revealed a preferential CD4^+^ T cell enrichment in TLS-like structures (B3-type lesions), and T cell-dominant aggregates (T-type lesions). In contrast, while abundant in M and T type lesions, the border of necrotic lesions (N) had the most CD8^+^ T cells relative to other lesions. Focusing on B3 type lesion, additional multiplex immunohistochemistry staining for BCL6 and PNAd confirmed the presence of TLS with germinal-center and high endothelial venules (HEVs) in B3 type lesions (Fig. 4B). Spatial association analysis among B3-type lesions (*n* = 110) revealed that as expected Tfh cells (C10) were positioned closer to the centralized B cells than proinflammatory CD4^+^ T cells (C08) in TLS (Fig. 4C). The localization preference analysis further confirmed that most Tfh cells presented in the center of TLS-like B3-type lesions, while proinflammatory CD4^+^ T cells were broadly dispersed, and proinflammatory CD8^+^ T cells were positioned further toward the periphery (Fig. 4D and 4E). These findings uncover the structured organization of distinct effector T cell populations within TLS in TB affected Lungs.

**Fig. 4.**
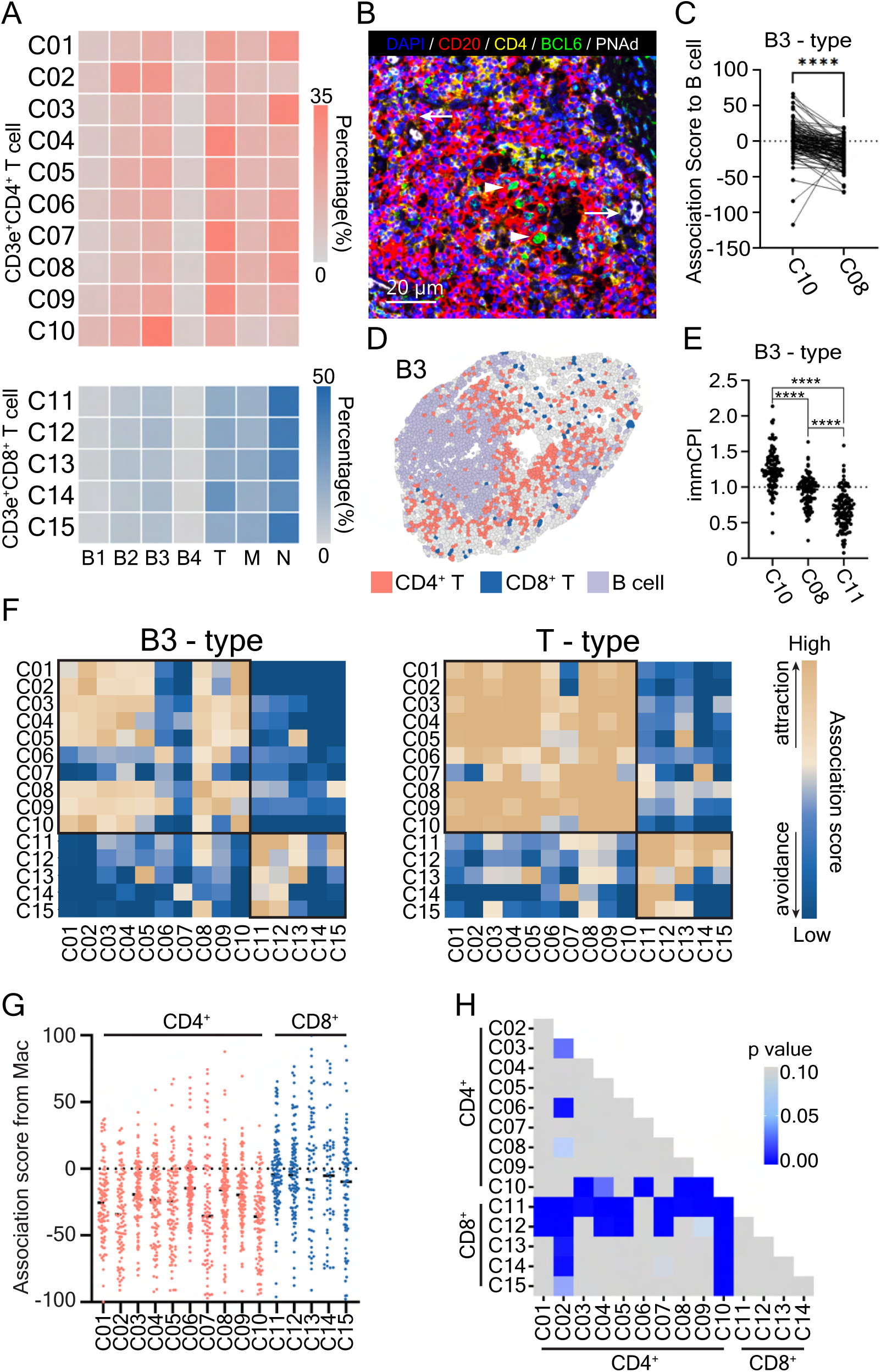
CD4^+^ and CD8^+^ T cells are spatially segregated and display different relationships with other immune cell populations. **(A)** Heatmap showing the relative abundance of CD4^+^ (top) and CD8^+^ (bottom) T cell sub-populations in each lesion type. Rows are ordered based on the similarity in marker expression patterns. N, the border of necrotic lesions. **(B)** Representative OPAL multi-plex image illustrating a germinal center-like structure in a B3-type lesion. The image with merged DAPI, CD20, BCL6, and PNAd is shown. Arrows indicate PNAd^+^ high endothelial venules and arrow heads indicate BCL6^+^ B cells. **(C)** Paired analysis comparing the spatial association relationships between B cells and Tfh (C10) or proinflammatory CD4^+^ T cells (C08) in all B3-type lesions. Each line indicates a lesion. The level of statistical significance is determined using Wilcoxon signed-rank test, **** *P* < 0.0001. **(D)** Cell type pheno-map of a representative B3-type lesion with cell type indicated. **(E)** Scatter plots showing the spatial centrality index (immCPI) of Tfh (C10), proinflammatory CD4^+^ T cells (C08), and proinflammatory CD8^+^ T cells (C11) in all B3-type lesions (*n* = 110). The level of statistical significance is determined using a Mann-Whitney U test, **** *P* < 0.0001. **(F)** Heatmap showing the median value of pair-wised spatial association relationships between CD4^+^ and CD8^+^ T cell subsets in B3 (left) and T (right) type lesions. **(G)** Column scatter plot showing spatial relationship between macrophages and CD4^+^ and CD8^+^ T cell subtypes in T-type lesions. **(H)** Heatmap indicating statistical significance from post hoc multi-group comparison results in (G) using a One-way ANOVA test, the level of significance was set at *P* = 0.05.

To investigate the spatial relationship among CD4^+^ and CD8^+^ T cell subsets within B3-type lesions, we performed pairwise spatial association analyses of their respective T cell clusters. We found that CD4⁺ and CD8⁺ T cell subsets preferentially co-localize with cells of the same lineage, suggesting that these two T cell lineages occupy distinct spatial niches within the tissue (Fig. 4F, left panel). This segregation was even more pronounced in T-type lesions (Fig. 4F, right panel).

We next assessed spatial associations between T cells and macrophages in T-type lesions characterized by abundant T cell and macrophage populations. Pairwise analyses revealed that most CD4^+^ T lymphocytes and approximately half of the CD8^+^ T cells within T-type lesions exhibited a clear avoidance from macrophages (Fig. 4G and 4H). The proinflammatory CD8^+^ T cells (C11) and progenitor-like CD8^+^ T cells (C12) had a significantly stronger spatial association with macrophages than their CD4^+^ counterparts (Fig. 4G and 4H). These observations reveal that CD4⁺ and CD8⁺ T cells occupy distinct spatial niches, with CD8⁺ T cells positioned closer to macrophages than their CD4⁺ counterparts, suggesting that CD8⁺ T cells may play a more prominent role in driving local inflammation in certain lesions.

### The border of necrotizing lesion harbors diverse sub-lesion microenvironments

We next examined immune cell organization in caseous lesions, representing a more damaging and advanced TB pathology. The lesions are characterized by a large hollow core containing largely dead and dying cells surrounded by a dense cellular structure, walling off the core region from neighboring area (*14*) (Fig. 5A). Historically, examining entire necrotizing lesions has been challenging because of their large size. We have previously shown that in necrotizing lesions CD14^+^CD68^+^ macrophage /monocyte populations preferentially localize at the inner necrotizing border (*9*). However, we have also observed an additional level of diversity in the cellular composition depending on the position that is examined around the circumference of necrotizing lesions. We observed that the expression of some cellular markers was polarized towards one side of necrotizing lesions compared to the other side (Fig. 5A).

**Fig. 5.**
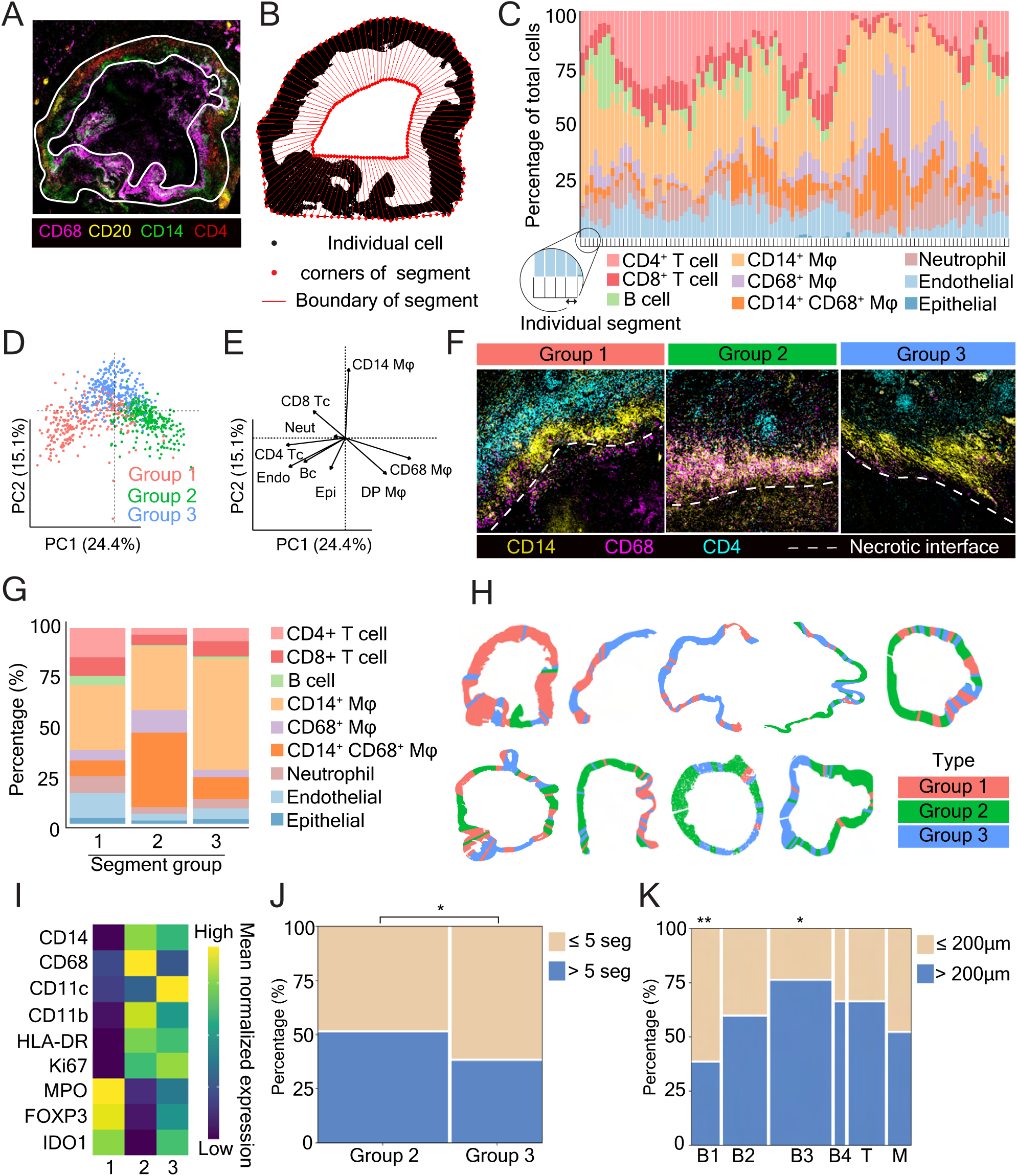
Cellular structure and immune response vary along the circumference of necrotizing lesions. **(A)** Composite immunofluorescence image of a necrotizing lesion with selected major immune markers shown. **(B)** Schematic diagram of the division of a lesion border into 100 individual segments each comprising approximately 1% of the lesion. **(C)** Percentage of each cell type across 100 segments from the lesion in (A) in order of their place around the lesion circumference. Each color corresponds to the indicated cell type. **(D)** PCA plot visualization of all segments across 9 necrotizing lesions. Each symbol represents an individual segment, and each color corresponds to one segment group. **(E)** PCA loading plot representing the contributing variables to the PCA plot in (D). DP Mφ, CD14^+^CD68^+^ Mφ. **(F)** Representative composite immunofluorescence images of necrotizing lesion areas of each segment group indicated in (D). **(G)** Mean percentage of each cell type in each segment group. Bars are colored according to indicated cell types. **(H)** Diagram of distribution of 3 segment groups along necrotizing circumference for each necrotizing lesion. Each color represents a segment group indicated in (D). **(I)** Heatmap of normalized expression of selected tissue markers (row) against segment group (column). **(J)** Mosaic plot showing proportion of group 2 and 3 segment localized within and outside 5 segment distance from the closest group 1 segment. **(K)** Mosaic plot showing proportion of lesion types localized within and outside 200 µm range to the closest border of necrotic granuloma. The level of statistical significance in (I) and (J) are determined using a Chi-square test, ** *P* < 0.01, * *P* < 0.05.

To quantify the localisation of each cell type around the circumference of necrotizing lesions, we divided the border of each necrotizing granuloma into 100 individual sections (each comprising 1% of the circumference) (Fig. 5B). This allowed us to visualize changes in the proportion of each cell-type present along the lesion border (Fig. 5C). This analysis was performed across nine necrotizing lesions from six patients. We pooled the segments from all lesions and performed PCA on the proportions of cell types within each segment to characterize the cellular diversity along the lesion border. We found that the lesions could be divided into 3 groups (Fig. 5D – 5G). Segment group 1 was the most enriched in CD14^+^CD68^-^ macrophages, CD4^+^ and CD8^+^ T cells, CD31^+^ endothelial cells, and CD15^+^MPO^+^ neutrophils. Segment group 2 was defined by its enrichment of CD14^+^CD68^+^ macrophages whereas segment group 3 was defined by its enrichment of CD14^+^CD68^-^ macrophages. We observed that all necrotizing lesions were comprised of multiple segment groups around their circumference, often intermingled with each other (Fig. 5H). Of note, the proportion of B cells was consistently found to be very low regardless of the segment group.

Given that segment group 1 contained high frequency of not only CD14^+^CD68^-^ macrophages but also T cells, we examined whether this was correlated to enhanced immune activity compared to other segment groups. We observed that the expression of MPO, IDO1 and FOXP3 was indeed largely confined to segment group 1 and to less extent to group 3, with minimal expression observed in group 2 segments (Fig. 5I). The elevated immune activity in group 3 segments appeared to be associated with heightened cellular interactions between group 1 and 3, since that group 3 segments were more frequently located within a five-segment distance to the immune-active group 1 segments compared to group 2 (*P* = 0.0204, Chi-square test) (Fig. 5J).

Finally, we investigated potential spatial relationships between the necrotic lesion border and surrounding 6 types of non-necrotic lesions, which were distributed in a satellite-like arrangement around the central cavity. Consistent with our previous findings (*9*), B cell aggregates (B1 type lesions) were significantly enriched within 200 μm of the necrotic lesions (*P* = 0.0082, Chi-square test) (Fig. 5K). In contrast, TLS (B3 type lesions) exhibited a spatial avoidance of necrotic lesions (*P* = 0.0126, Chi-square test). The rest of the lesion types did not display a consistent spatial trend relative to the necrotic border, revealing a lesion-type dependent spatial relationship between necrotizing and non-necrotizing lesions.

### Hierarchical spatial analysis reveals cellular architecture associated with distinct clinical manifestations

From tissue-wide quantification to cellular analysis, our hierarchical spatial assessment revealed that non-necrotizing lesions exhibit diverse and dynamic cellular compositions embedded with distinct microenvironments. To link these spatial features to clinical outcomes, we performed comparative analysis of samples from patients with clinical and subclinical TB (Fig. 6A). At the tissue-wide level, overall cellular and community compositions did not differ significantly between the two groups. However, at lesion level, patients with clinical TB (*n* = 5) exhibited significantly fewer M-type lesions (*P* = 0.0079, Mann-Whitney U test) compared to those with subclinical TB (Fig. 6B and 6C). Similarly, clinical samples contained a higher number of intact alveolar inflammation B4-type lesions (*P* = 0.0476, Mann-Whitney U test), suggesting increased and progressing tissue inflammation (fig. S4A). Given our finding of macrophage and B cell segregation across lesion types (Fig. 3H), it is unsurprising to observe the frequency of B cell enriched lesions was trending higher in the clinical patient samples than subclinical samples (Fig. S4A).

**Fig. 6.**
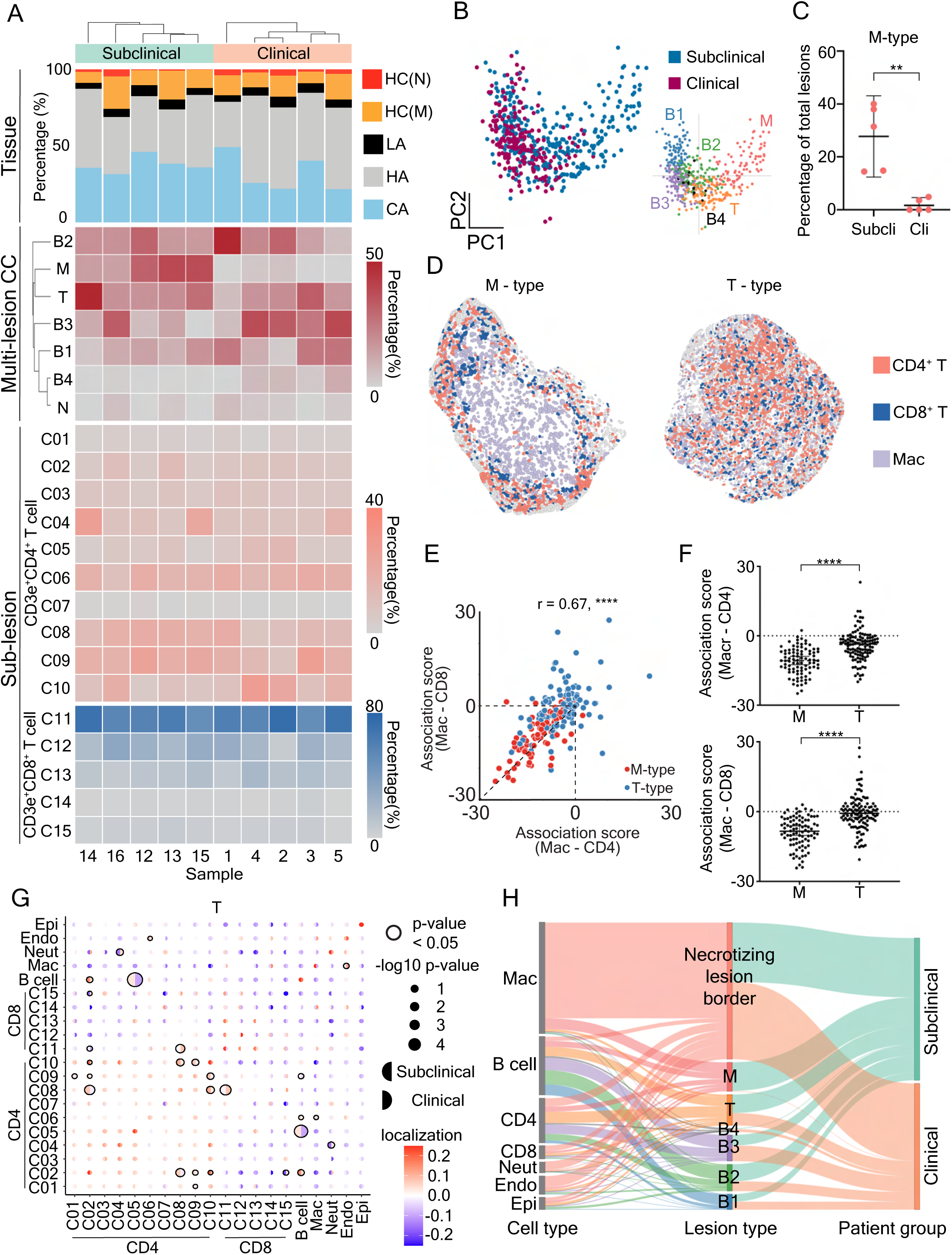
Hierarchical spatial analysis reveals cellular architecture associated with distinct clinical manifestations. **(A)** Heatmap showing the clinical group, CC percentage, lesion type percentage, CD4^+^ and CD8^+^ T cell sub-populations percentage of human TB lung tissues (*n* = 10). HC(N), histiocytic cuff (neutrophil enriched); HC(M), histiocytic cuff (macrophage enriched); LA, lymphoid aggregates; HA, healthy alveoli; CA, consolidated area. **(B)** PCA plot of the distribution of the lesions from clinical and subclinical TB patients (n = 525). The lesions are colored based on clinical symptom status (left), and lesion type (right). **(C)** Percentage of M-type lesions among total lesions in each patient of the clinical or subclinical TB group. Each symbol represents a patient. The statistical significance between two groups is determined using a Mann-Whitney U test, ** *P* < 0.01. **(D)** Cell type pheno-map of representative M and T-type lesions with cell type as indicated. **(E)** Scatter plot showing association scores between macrophages and CD4^+^ versus CD8^+^ T across M and T-type lesions. Correlation is determined using Spearman’s correlation test, **** *P* < 0.0001. **(F)** Scatter plot comparing the spatial association scores from macrophage to CD4 (top) or CD8 (bottom) between M and T-type lesions. Each dot indicates a lesion. The level of statistical significance is determined using Mann-Whitney U test, **** *P* < 0.0001. **(G)** Heatmap showing pair-wise comparison of spatial association scores between subclinical and clinical groups. The proximity of each cell population to other populations indicated in T-type lesions were determined using 50 μm radius range. **(H)** Alluvial diagram highlighting the connections between in-lesion, cross lesion, and clinical cohorts. Left represents the major cell types in lesions, middle shows stratified lesion types, right shows the clinical cohorts. Line thickness represents the proportion within each group.

M-type lesions, which were more common in subclinical group, are characterized by dense myeloid cores, macrophages were generally spatially segregated from both CD4^+^ and CD8^+^T cells. In contrast, T-type lesions, another T cell-enrich structure, exhibited a relatively more diffuse arrangement, with some T cells and macrophages in closer spatial proximity (Fig. 6D). A positive correlation between CD4^+^ and CD8^+^ T cell proximity to macrophages (*r* = 0.66, *P* < 0.0001) was observed, indicating similar spatial behavior of both T cell populations relative to macrophages (Fig. 6E). M-type lesions displayed spatial avoidance between T cells and macrophages, while T-type lesions, characterized by high T cell infiltration, showed significantly closer associations, with some lesions demonstrating positive interaction scores (Fig. 6F).

Further dissection of T-type lesions revealed differential spatial organization of CD4^+^ T cell subsets between two clinical groups. In subclinical TB, proinflammatory CD4^+^ T cells (C08), progenitor CD4^+^ T cells (C09), and Tfh cells (C10) were co-localized (Fig. 6G). In contrast, these spatial associations were significantly reduced in the clinical TB group, potentially reflecting weakened local immune responses. A similar trend was observed in TLS-like B3-type lesions, where activated CD4^+^ T cells (C02) and proliferating CD4 ^+^ T cells (C01), as well as ICOS^+^ population(C05) and Tfh cells (C10), showed higher spatial association in subclinical samples (fig. S4B). Overall, these data suggest that a weakened organization of T cell subpopulations within lesions is associated with an increased local immune response in clinical TB.

To integrate multi-scale spatial data with clinical observations, an alluvial diagram was generated (Fig. 6H). It revealed that macrophages were the most abundant immune cells in TB lungs, primarily localizing to the borders of necrotizing lesions, with a smaller fraction found in M-type lesions. While necrotizing lesions, indicative of more destructive pathology, were present in both clinical and subclinical TB, M-type lesions, suggestive of milder pathology, were mainly observed in subclinical cases. This distribution implies that macrophages may adopt context-dependent roles, contributing either to tissue damage or immune regulation.

## DISCUSSION

Human TB is characterized by a wide array of tissue pathologies that can differ significantly among individuals and even within the same patient. How the immune responses underlying the diverse tissue alterations are regulated to minimize local inflammation and preserve normal pulmonary function remains incompletely understood. In this study, we analyzed over 500 lesions from the lungs of patients with either clinical or subclinical TB and revealed that, while all major immune cell types were present in TB-affected lung tissue, many of these cells were spatially segregated from one another both across and within TB lesions. We propose that spatial sequestration of macrophages from effector lymphocytes represent a mechanism aimed at controlling excessive inflammatory responses in diseased lung tissues. Consistent with this notion, we observed that, lesions with prominent spatial separation between T cells and macrophages were significantly less common in clinical TB patients compared to those with subclinical disease.

Our research has revealed a previously unrecognized function of B cells in human pulmonary TB through their interaction with macrophages. While previous studies have reported cross-regulation between macrophages and B cells with varying outcomes in other disease models (*32–34*), we showed that the competition between the effector cell populations correlates with the variations of non-necrotizing lesion architectures in tissue. This finding aligns with our previous report (*9*) and a spatial transcriptomics study that showed a division of myeloid and lymphoid gene responses in human TB lung tissue (*35*). However, it is worth noting that organization of the distinct microenvironments in necrotizing lesion appears to be independent of B lymphocytes given their low abundance. Understanding the mechanisms dictating the mutual exclusivity between B cells and macrophages may offer insights into potential strategies for managing lung pathology.

Little is known about the immune response associated with different stages of human TB (*12, 36*). This study shows that while full spectrum of lesions was presented in the subclinical TB stage, M-type lesions were less common in the active stage of the disease. The spatial organization of immune cells in M-type lesions resembles the stereotypical “granuloma” structure described in the literature. The unique cellular organization, avoidance between T cells and macrophages (the scarcity of T cells in the macrophage core), could play a lesion-dependent role in the early inflammatory processes. The range of cytokine communication has been shown to be between 30-150 microns depending on the density of cytokine-receiving cells (*37*). It is possible that some macrophages in the core region of large M-type lesions are not effectively stimulated by cytokines produced by the T cells due to the distance between these cells. The reduced communication between T cells and macrophages in certain lesions may impair innate cell activation, compromising bacterial control while preventing excessive tissue damage. Interestingly, a high abundance of macrophages is found in the inflammatory necrotizing border and contained, M-type non-necrotizing lesions, suggesting macrophage populations may play distinct role in the two types of inflammatory foci. Understanding the role of macrophages in various lesion types and clinical stages opens avenues for potential therapeutic interventions aimed at modulating macrophage activity.

The preferential accumulation of various T cell subsets within certain TB lesions suggests that these inflammatory foci may play different roles in recruiting and/or maintaining T cell effector or selective survival functions in TB-infected lungs. In B3 lesions, follicular helper T cells were positioned close to germinal center-like structures containing proliferating BCL6^+^CD20^+^ B cells, indicating an intimate interaction between follicular helper T cells and B cells and hinting at the development of TLS that support T-B cell interactions. The formation of TLS in TB-affected lungs has been reported in Mtb-infected humans, non-human primates, and mice (*38*). TLS supports the regional differentiation and stabilization of effector T cells in autoimmune disease (*39*) and cancers (*40, 41*) in humans. A similar mechanism may operate in the lungs of TB patients to sustain or regulate the regional response to Mtb infection. This hypothesis is supported by the finding that B3-type lesions serve as preferred sites for CD4^+^ T cell accumulation, housing diverse effector cell populations and stem-like TCF1^+^CD4^+^ T cells. Interestingly, their frequency diminishes significantly in the highly inflamed borders of necrotizing regions. In mice, these TCF1^+^CD4^+^ T cells sustain the development of effector cells (*27, 42*), suggesting a possibility that the CD4^+^ T cell population in B3-type may serve as a reservoir to replenish effector T cell pool locally in the tissue.

In B3 and T-type lesions, we found that CD4^+^ and CD8^+^ T cells tend to occupy distinct niches within the same lesion. This segregation pattern is similar to the distribution of T cells in naive lymph nodes, where CD4^+^ T cells are predominantly located at the periphery of the paracortex, while CD8^+^ T cells are more centrally positioned (*43*). The reasons behind this spatial segregation in TB lesions remain unclear, although this could result from different chemokine niches presented in the lesions. Interestingly, our data suggest that CD8^+^ T cells, particularly the GZMA and GZMK expressing subset, are often found closer to macrophages than CD4^+^ T cells, highlighting the potential role of CD8^+^ T cells in interacting with macrophages in TB. The function of GZMA and GZMK-expressing CD4^+^ T cells in humans is still not well understood. GZMK expression in T cells is influenced by CD15^+^ neutrophils (*44*) and has recently been implicated in complement activation (*29, 30*). Additionally, our earlier research on TB pleural effusions showed that, unlike GZMB^+^ T cells which are predominantly found in the bloodstream, GZMK^+^ T cells are more prevalent at the infection site in the pleural cavity (*45*). Collectively, these findings suggest that GZMK-expressing T cells play a significant role in regulating tissue inflammation.

Limitations of the study include the relatively low-plex nature of current spatial proteomics compared to single-cell RNA sequencing, and the restriction of results to tissue sites collected during surgery. Additionally, the sample size of this study is small. Future work is required to validate the reported findings. Finally, all patients included here were immunized with BCG and some of these patients have also been treated with anti-TB drugs. It remains to be established whether and how vaccination and chemotherapy modulate the regional immune response. Nonetheless, the information reported here may be useful for the development of adjunct host-directed therapies for multi or total drug-resistant TB.

## MATERIALS AND METHODS

### Clinical information

This study utilized a retrospective cohort of patients from the Shenzhen Third People’s Hospital. The study was approved by the Research Ethics Committees of the Shenzhen Third People’s Hospital (2020–019) and Sydney Local Health District (X18-0367). All archival specimens were analysed with no active participation of human subjects.

### FFPE sample collection and processing

Formalin-fixed, paraffin-embedded (FFPE) samples were obtained from consenting patients diagnosed or suspected of having TB in China (table S1). To explore granuloma heterogeneity and its potential relevance to disease progression, we examined specimens from two infection states. The clinical TB group included lung tissues from patients with progressive TB undergoing resection after partial or failed response to anti-TB treatment (table S1, patient ID 1-11), while a second set consisted of patients with incidental or subclinical lung nodules that were ultimately confirmed as TB by histopathological examination post-resection (table S1, patient ID 12-22). Each section was reviewed by two individual clinical pathologists to confirm Mtb infection.

For slide preparation, FFPE blocks of TB lung samples were sectioned by microtome (Leica, #14051756235) at 4 μm thick located in an 18 mm × 35 mm imaging area on Super Frost Menzel positive charged microscope slides (Epredia, HL26766). The sections were baked at 55 ℃ for 1 hour and stored at 4 ℃ cold room before further staining.

### Antibody panel design and validation for PCF

A panel of 35 markers (fig. S1A, table S2) was designed to target immune and stromal cells. The markers were chosen based on prior panels for multiplexed immunofluorescent studies (*9*), and our scRNA-seq data (*45*). Primary antibodies were chosen for testing which had FFPE validation by the vendors and clinical pathologists. Primary antibodies for custom conjugation and clones were reviewed by experienced immunologists. The selected antibodies were validated with chromogenic IHC staining in serial sections of tonsil and human TB lung tissues.

### PhenoCycler-Fusion antibody conjugation

Akoya antibodies were purchased pre-conjugated to their respective PCF barcodes (table S2). All other antibodies were custom conjugated to their respective PCF barcodes according to Akoya’s PhenoCycler-Fusion user guide using the antibody conjugation kit (Akoya Biosciences, Cat# 7000009). Briefly, 50 μg of carrier-free antibodies were concentrated by centrifugation in 50 kDa MWCO filters (EMD Millipore, Cat# UFC505096) and incubated in the antibody disulfide reduction master mix for 30 minutes. After buffer exchange of the antibodies to conjugation solution by centrifugation, addition of conjugation solution and centrifugation, respective PCF barcodes resuspended in conjugation solution were added to the concentrated antibody and incubated for 2 hours at room temperature. Conjugated antibodies were purified by 3 buffer exchanges with purification solution. Finally, 100 μL of antibody storage buffer was added to the concentrated purified antibodies. Custom conjugated antibodies were titrated according to the manufacturer’s instructions (Akoya Biosciences), based on a validation imaging run using control tissue matched to the tissue type.

### PhenoCycler Fusion sample processing and staining

FFPE sections were baked at 65 ℃ for 2 hours and dewaxed through xylene for 10 minutes. Gradients of ethanol and deionized purified water were used for dehydration (100% ethanol for 10 mins, 90% ethanol for 5 mins, 70% ethanol for 5 mins, 50% ethanol for 5 mins, 30% ethanol for 5 mins, ddH2O for 10 mins). Antigen retrieval was performed by pressure cooker at 120 ℃ for 20 minutes in pH 9 Antigen retrieval buffer (Akoya, #AR9001KT). Tissues were incubated in Hydration buffer (Akoya, #7000008) for 2 × 2 minutes and Staining buffer (Akoya, #7000008) for 30 minutes. Tissues were stained under room temperature with 200μL mixture of primary conjugated antibodies based on the optimized panel for 3 hours.

The tissues were washed in Staining buffer (Akoya, #7000008) for two 2-minute washing step and then fixed in post-staining solution (1.6% formaldehyde in PBS) for 10 minutes. Tissues were then washed in PBS for 3 × 1 minute and then incubated in ice-cold methanol for 5 minutes. Tissues were washed in PBS for 3 × 1minute and fixed by fixing reagent (Akoya, #7000008) for 20 minutes. The fixed tissues were washed in PBS for 3 × 1 minute and then ready for Flow cell assembly and multiplexed imaging in PhenoCycler-Fusion (Akoya Bioscience).

### PhenoCycler-Fusion sample imaging

A 96-well plate of reporter stains with Nuclear Stain (Akoya, #7000003) was prepared according to manufacturer’s instructions (Akoya Biosciences). Stained slides were loaded onto the Fusion Stage Insert and the Reporter Plate was loaded into the PhenoCycler Microfluidics Instrument. Images were acquired under the high sensitivity settings (8-bit) in the PhenoCycler Fusion software, with the exposure time for DAPI set to 2 milliseconds. Slides were imaged in the fluorescence channels followed by a gentle wash to rehybridize and remove reporters. Raw .tiff image files were acquired after each cycle, and final .qptiff image was processed by Fusion software (1.0.3 or 1.0.5).

### Cell segmentation and quality control

Acquired images were registered into QuPath (0.4.3) (*46*). Nuclear segmentation was first performed using StarDist (*19*) with pretrained model “2D_versatile_fluo” applied to the DAPI channel. The segmentation was performed at the resolution 0.5 µm per pixel aligned to the acquired image dataset. Cytoplasm segmentation was then estimated from nuclear expansion by morphological dilation (5 µm). Quality control of the data was performed qualitatively on each individual marker channel and filtering was conducted to exclude markers and tissue regions with inadequate quality. The single-cell protein measurements were obtained as the mean fluorescent intensity (MFI) per marker within the cell mask, and the centroid of each cell was defined by the x-y coordinates in the images. Cells were manually removed if they fall in regions of tissue folding or detachment, or in areas with noticeable artifactual staining. The finalized MFI table was exported as .csv for downstream processing and analysis.

### Unsupervised clustering and cell phenotyping

Cell clustering was performed using FuseSOM (*20*). Z-score normalization was applied across all cells for each marker such that each marker has a mean equal to 0 and a standard deviation equals to 1. The normalized MFI expression table for all markers and all cells from each image was used for unsupervised clustering. The cells were over-clustered using K-means algorithms with cell lineage markers (CD45, CD3e, CD20, CD4, CD8, CD15, MPO, CD14, CD68, PAN-CK, CD31). Clusters were manually annotated into major cell lineages (B cell, CD4 Tc, CD8 Tc, DP Mac, CD14 Mac, CD68 Mac, Neutrophils, Epithelial, and Endothelial cells). Clusters with similar expression patterns were combined into one phenotype, and clusters with false expression pattern were removed after inspected on raw immunofluorescence images.

### Cellular communities and spatial analysis

The spatial interaction between different cell phenotypes were quantified using LisaClust (*21*). This method provides critical information about the relative co-occurrence of cell types within a cell’s local environment. In this study, a cell’s community was first defined using PCA analysis, where for each cell, the cell-type percentages in its neighboring area were calculated. This step produces a matrix where cells are characterized by the percentages of cell types in their local neighborhood. K-means clustering (*K* = 5) was performed on all cells based on these percentage within 50μm radius. The optimal number of cellular communities was determined as 5 because it presents sufficient communities that separate the cells into small groups differentiated by phenotype variability over the technical variability.

### TB pathology segmentation model development and validation

A “human-in-the-loop” pipeline was developed to train a TB pathology segmentation model capable of tissue characterization with near human-level performance. Out of 22 post-PCF H&E-stained tissue sections, 7 were partially annotated by human pathologists and used to train a convolutional neural network (CNN) with an encoder–decoder architecture derived from the U-Net model. In this architecture, the encoder learns hierarchical features from 256 × 256-pixel patches, while the decoder reconstructs the segmentation output. Model optimization was performed using a standard cross-entropy loss function. Training was conducted on NVIDIA RTX A6000 GPUs for 50 epochs, with a batch size of 8, a learning rate of 1 × 10^-4^, and the Adam optimizer.

The trained model was then evaluated on the remaining H&E-stained tissue sections. Predicted segmentation masks were compared to annotations of cellular communities using Pearson’s correlation, providing a measure of the linear relationship between model output and the proteomic cellular community analysis results.

### Delineation of necrotizing lesions into segments

The borders of necrotizing lesions were delineated visually using both H&E and PCF images to determine the edges of the dense immune infiltrate surrounding necrotic cavities. The .csv file corresponding to the cells from each necrotizing lesion was imported into R with the cell identities previously determined. A line equivalent to the border of each lesion was generated using the concaveman() function from the Concaveman package. The degree of concavity was selected to follow the general contours of the lesion while also low enough so that any indents in the lesion would be smoothed over. The resample_polyline() function was then applied to the lesion border to generate points along the line, with the interval length selected for each lesion such that exactly 100 evenly spaced points would be generated. The st_buffer function from the sf package (*47*) was then used to replicate a smaller inner copy of the lesion border, contained entirely within the central necrotic area of each cavity, with 100 evenly spaced points also generated on this line as above. Each of the 100 lesion segments was then defined using 4 points (2 from outer border and 2 from inner) from the lesion border, and the point.in.polygon() function was used to assign each cell from the lesion border into 1 of the 100 segments.

### Analysis of necrotic lesion segments

The cellular composition of each segment was visualized with stacked bar plots generated from the geom_bar() function from the ggplot2 package (*48*). Lesion clustering analysis was performed using PCA with a density of CD8+, CD20+, CD4+, and CD68+ cells used as parameters. PCA was performed in R with the prcomp() function with “scale = True”. PCA results were visualized using the factoextra package with functions fviz_pca_ind() and fviz_pca_var(). The distance of segments from types 2 and 3 to the closest segment of type 1 was determined by assigning a number to each segment (from 1 to 100) and determining the absolute number of segments from types 2 and 3 to type 1. As the segments were arranged circularly around the lesion (with lesion 1 and 100 adjacent to each other), rather than linearly, an adjustment to the calculation was made such that the starting point of the lesion enumeration was changed for each segment such that the segment being measured was assigned to number 50, with other segments evenly divided around it. The heatmap of functional marker expression was generated using the heatmap() function in R.

### SAM lesion segmentation

The Segment Anything Model (SAM) (*22, 23*) extension in QuPath (0.4.3) (*46*) was used to segment lymphoid-enriched lesions. PCF datasets, which included leukocyte markers (CD3e, CD20, CD14, CD68) and DAPI for cell morphology, were provided to the large vision model for lesion detection and boundary annotation. The ‘vit_l’ variant of SAM was used in multi-mask mode, generating three potential masks for each region; the one with the highest quality score was kept. Lesions with a quality score below 0.5 or containing fewer than 200 cells were considered poorly structured or small and excluded from further analysis.

### Total central preference index and immune cell central preference index calculation

The distance of each cell from the lesion boundary was measured using “signed distance to annotations 2D” feature in QuPath (0.4.3) (*46*). Custom groovy scripts were generated as previously described (*9*) to determine the median cell distance from the boundary and calculate the lesion area occupied by the inner and outer 50% of cells. From these measurements, we derived the total central preference index (tCPI) as follows:

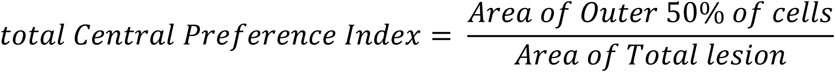

To compute the immune cell central preference index (immCPI), we applied the same distance measurements to subsets of cells defined by their immune markers. For each cell type, the immCPI was calculated by comparing the mean distance of those immune cells to the mean distance of all cells:

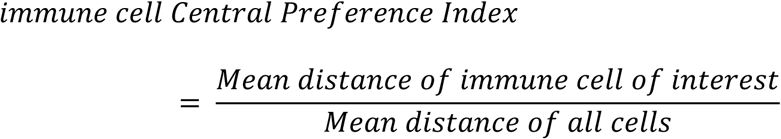

### Lesion cell clustering

Hierarchical sub-clustering was performed individually for CD4^+^ and CD8^+^ T cells using Scanpy (version 1.9.5) (*24*), starting from normalized MFI expression table of selected T cell (Ki-67, FOXP3, GZMA, GZMB, GZMK, ICOS, TCF1, PD-1, HLA-DR) derived from annotated T cells within extracted lesion area. To generate an over-clustered set of T cells, PCA was applied separately to CD4^+^ and CD8^+^ T cell subset, retaining the top 11 principal components (n_comp = 11). A k-nearest neighbor (k-NN) graph was constructed using 50 neighbors (n_neighbors = 50), and clustering was performed using the Leiden algorithm at a resolution of 0.5 (resolution = 0.5).

Differential expression analysis was conducted using Wilcoxon rank-sum test (method = ‘wilcoxon’), comparing clusters based on Leiden assignments (groupby = ‘leiden_0.5’). Each cluster was manually inspected in the raw immunofluorescence images (PCF datasets), and clusters with similar expression patterns were merged, while cells with inconsistent or spurious marker profiles were removed.

### Spatial association analysis

Spatial cell-cell association analysis was performed using SpicyR (*31*). Colocalization of cell types was assessed using Ripley’s K-function, which quantifies the scaled average abundance of surrounding cell types within a 50 µm radius of target cells. Pairwise spatial associations were evaluated for all annotated cell types within each individual lesion. A positive value indicates a relative spatial association, whereas a negative value signifies spatial avoidance compared to a random spatial distribution.

### Multiplexed immunohistochemistry staining

FFPE TB lung tissue sections were baked at 65 °C for overnight, then de-waxed and rehydrated in a series of xylene and ethanol gradients. Antigen retrieval was performed using a pressure cooker in pH 9 buffer (Akoya Biosciences) for 20 minutes. Next, tissue sections were incubated for 1 hour at room temperature with primary antibodies against CD20 (1:100, L26, Abcam), CD4 (1:100, EPR6855, Abcam), and overnight under 4°C with BCL6 (1:100, EPR11410-43, Abcam), and PNAd (1:200, MECA-79, Novus Biologicals). The primary antibodies were detected with OPAL Polymer HRP Ms+Rb (Akoya Biosciences) or Rat HRP-conjugated secondary antibody (Invitrogen, #31476) for 15 minutes, followed by a 10-minute incubation with Tyramide signal amplification and Opal Fluorophore (Akoya Biosciences). The slides were then heated to 100 °C to strip the antibody-HRP complexes, and this cycle of staining and stripping was repeated until all markers were labeled. Following the final round of staining, tissue sections were incubated with DAPI for 10 minutes and mounted using Prolong Gold (Thermo Fisher Scientific, #P36930). Images were acquired with the Fusion imaging system V2.0 (Akoya Biosciences).

### Statistical analysis

Where indicated, data were analyzed for statistical significance and reported as *P* values. Statistical analyses were conducted using Prism 10 (GraphPad, La Jolla, CA). Comparisons between multiple groups were performed using the Kruskal-Wallis test, followed by Dunn’s multiple comparison test. Pairwise comparisons between two groups were conducted using either the Wilcoxon signed-rank test (for paired data) or the Mann-Whitney U test (for unpaired data). Associations between categorical variables were evaluated using the chi-square (χ²) test. To evaluate relationships between variables, Pearson’s correlation test was used for assessing linear correlations, while Spearman’s rank correlation test was applied for nonparametric associations. The correlation coefficient (*r*) and corresponding *P* value were reported for all correlation analyses, with the strength of association interpreted as follows: weak (0.10–0.39), moderate (0.40–0.69), and strong (≥0.70). Statistical significance was defined as *P* < 0.05. Significance levels are denoted as follows: \**P* < 0.05, \*\**P* < 0.01, \*\*\**P* < 0.001, \*\*\*\**P* < 0.0001.

## Acknowledgements

We thank Drs. Erin McCaffrey and Lina Daniel for invaluable discussions and critical reading of the manuscript.

## Funding

National Institutes of Health grant (U01AI166309), National Health and Medical Research Council Ideas Grant (2029659), Shenzhen Medical Research Fund (No. A2304001), National Key Research and Development Program (No. 2022YFC2302901), Shenzhen University 2035 Program for Excellent Research (2023B005), Guangdong Provincial Science and Technology Program (2019B030301009).

## Author contributions

Conceptualization: WX, CGF, and XC.

Methodology: WX, AJS, SM, XB, QY, YG, CQ, JW, MP, and EP.

Investigation: WX, AJS, SM, XB, QY, YG, TF, YZ, YD, QY, GD, GD, LF, YC and WJB.

Funding acquisition: CGF, XC, JDE, DLB, and EP. Project administration: CGF and XC.

Supervision: CGF and XC.

Writing – original draft: WX and CGF.

Writing – review and editing: all authors

## Competing interests

Authors declare that they have no competing interests.

## Data and materials availability

All key data are available in the main text or the supplementary materials. Raw imaging data, and analysis codes will be made publicly available upon acceptance of the manuscript.

---

**Figure S1.**
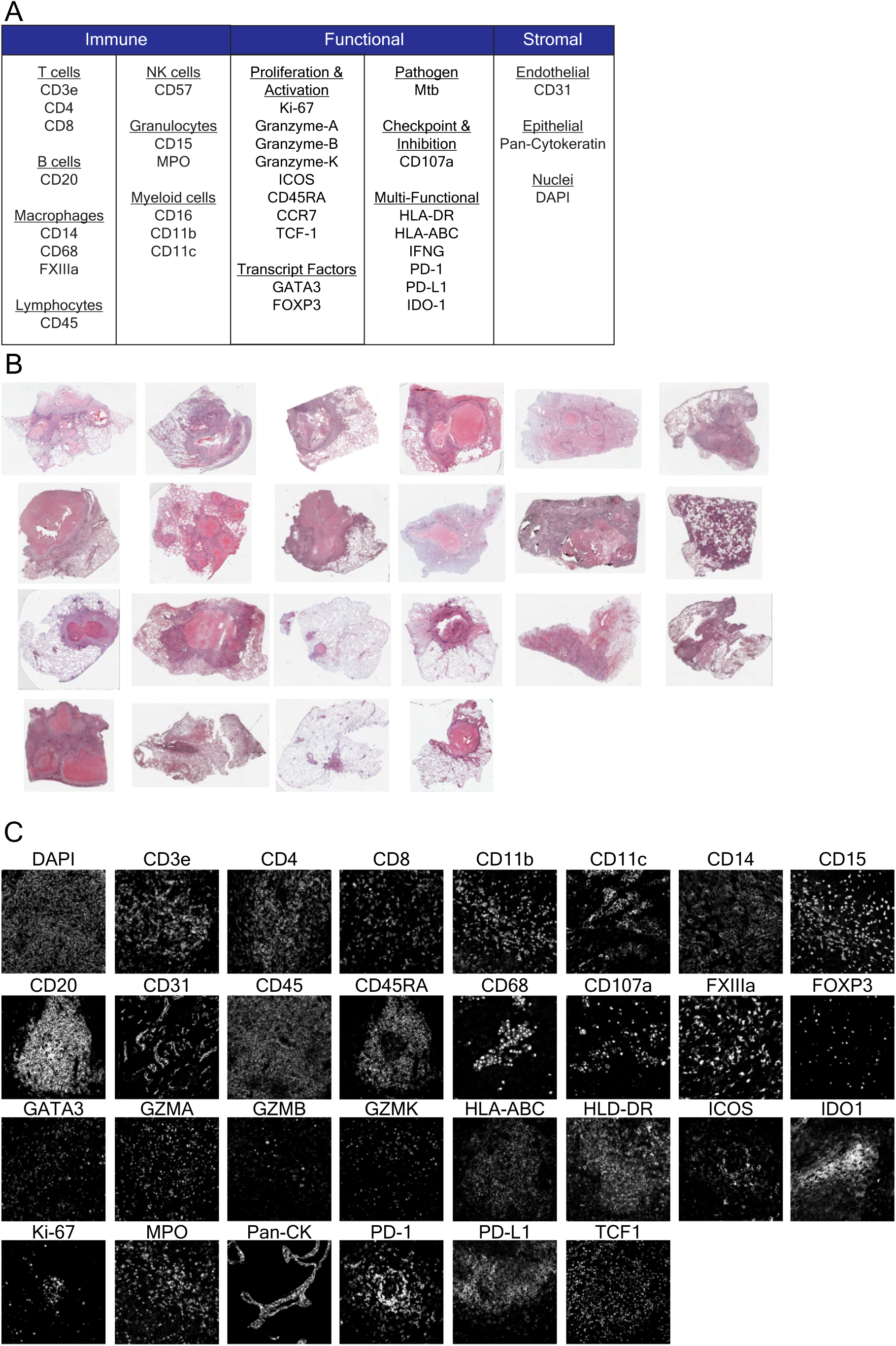
Multiplexed whole-slide immunofluorescence imaging of human TB lung tissue. **(A)** Multiplexed antibody panel grouped by marker category. **(B)** Hematoxylin and eosin-stained sections of samples post PCF imaging (n = 22). **(C)** Grayscale images of single-marker signal in PCF images.

**Figure S2.**
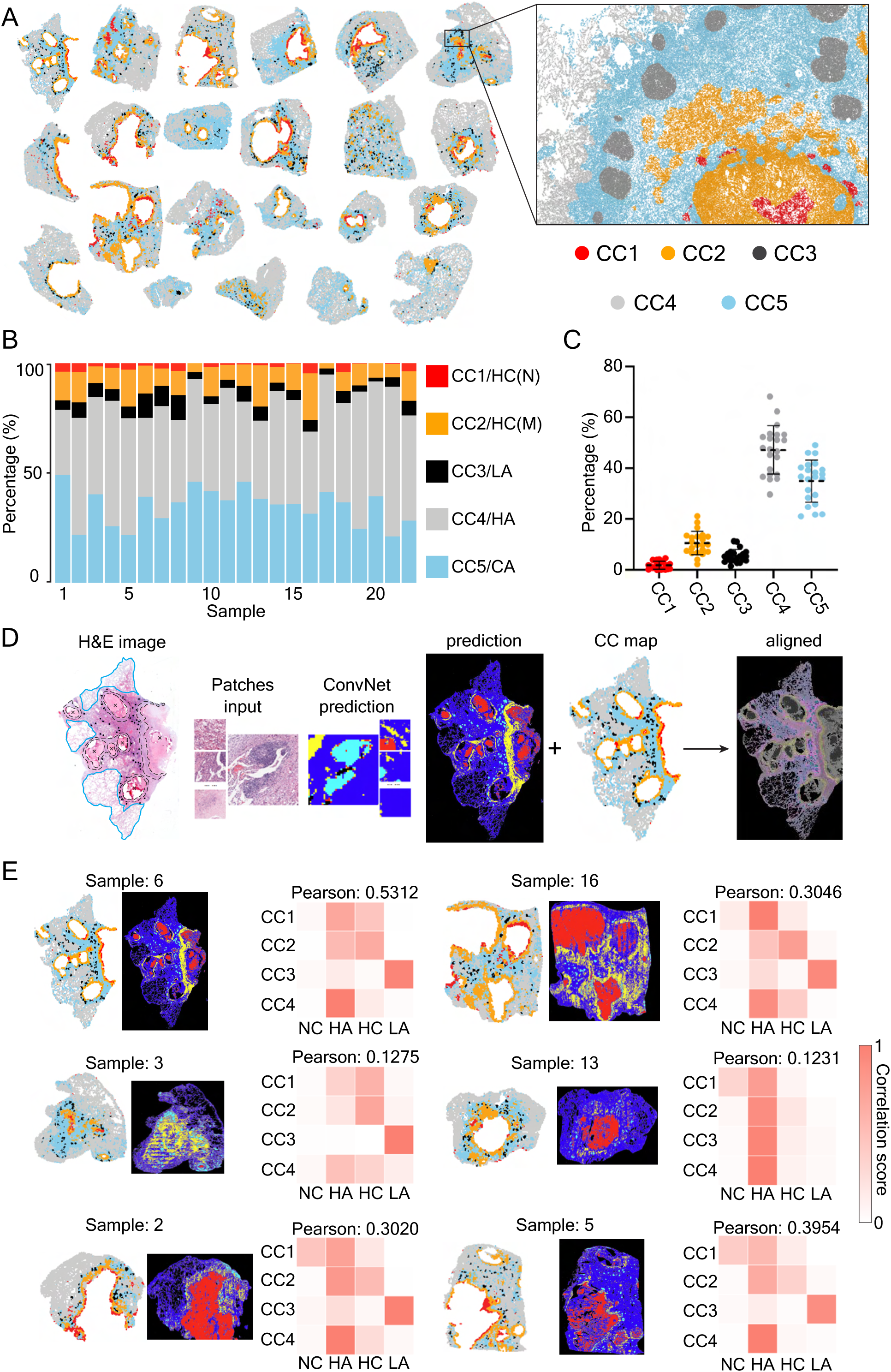
Cellular-community modeling of TB lung pathology. **(A)** CC phenotype maps of all samples (n = 22). **(B)** Percentage of defined CC among total segmented cells colored by CC type. Each column represents an individual clinical sample. **(C)** Scatter plot showing percentage of CC among all samples. Each symbol represents an individual clinical sample. **(D)** Schematic workflow for deep-learning-based pathology segmentation model training and correlation calculation against CC maps. **(E)** Correlation matrix of aligned samples (n = 6) showing correlation between defined CC and pathology prediction results. Each matrix indicated one clinical sample corresponding to the image on the left. HC(N), histiocytic cuff (neutrophil enriched); HC(M), histiocytic cuff (macrophage enriched); LA, lymphoid aggregates; HA, healthy alveoli; CA, consolidated area; NC, necrosis.

**Figure S3.**
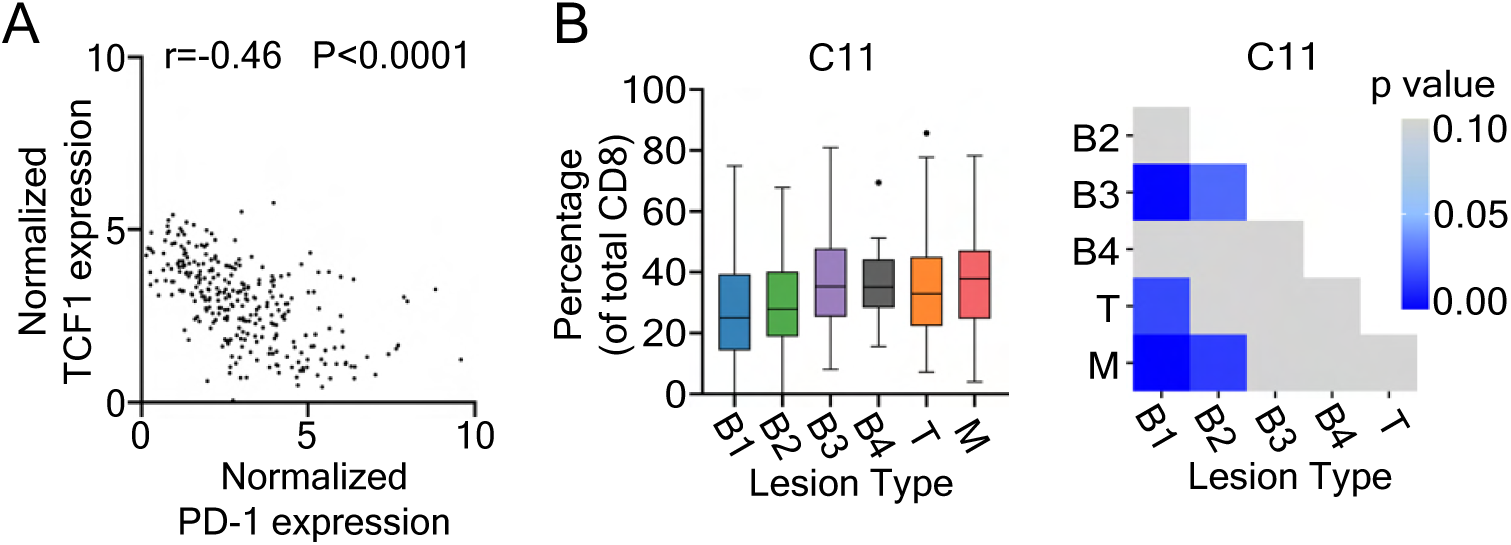
Immunoregulatory protein expression in Tfh cells and CD8 CTL localization across lesion types. **(A)** Linear relationship between normalized TCF1 expression levels and normalized PD-1 expression levels in B3-type lesions. Each dot represents one lesion. Correlation is determined using Spearman’s correlation test. **(B)** Box plot (left) showing the percentage of C11 among total CD8+ T cells in all lesion types, and heatmap (right) indicating statistical significance of multi-group comparison for each respective cluster with p value shown. The level of statistical significance is determined using One-way ANOVA, the level of significance was set at p = 0.05.

**Figure S4.**
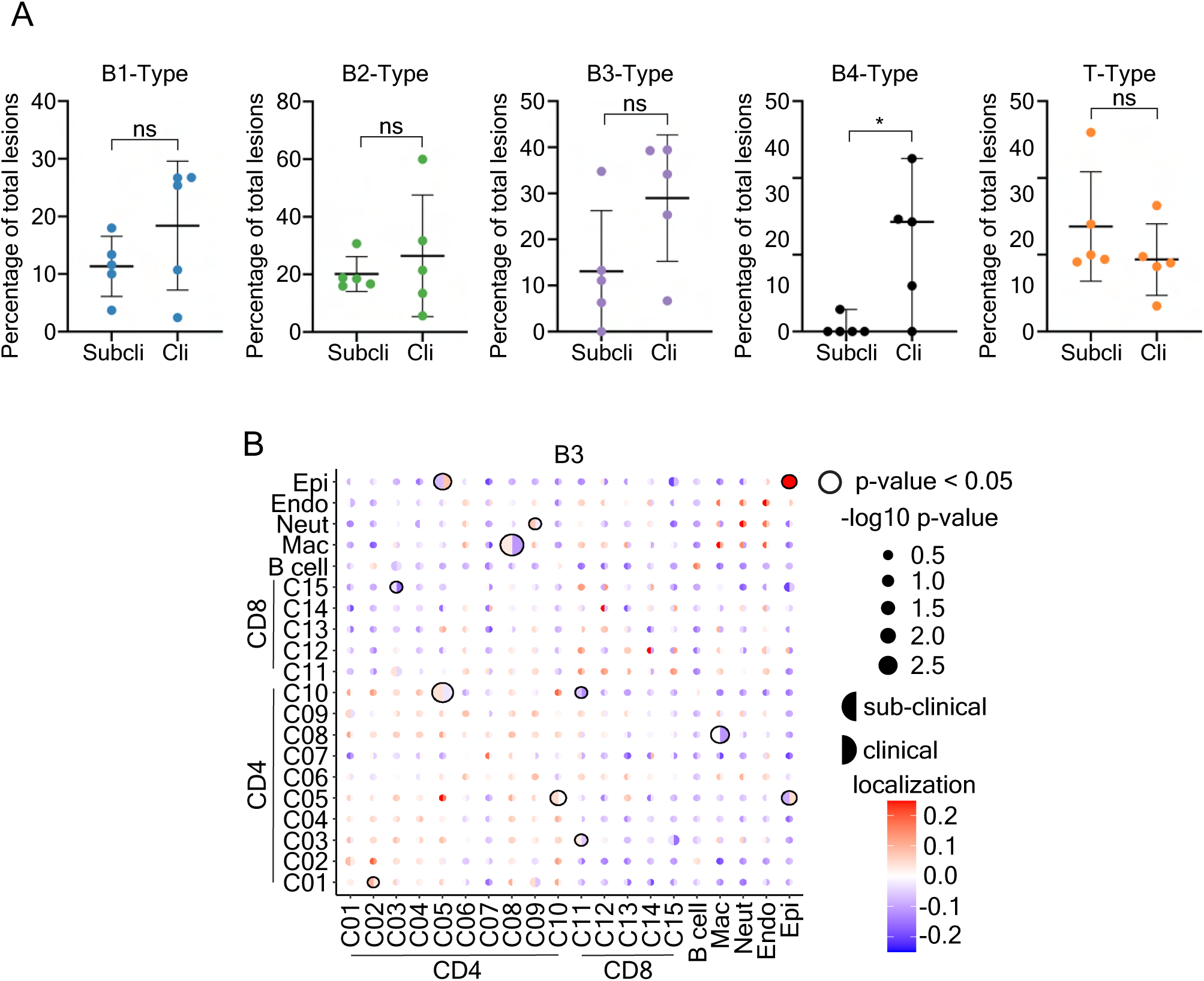
Spatial analysis reveals cellular organization in various type of lesions associated with distinct clinical manifestations. **(A)** Percentage of B1, B2, B3, B4, and T-type lesions among total lesions in each patient of the clinical or sub-clinical TB group. Each symbol represents a patient. The statistical significance between two groups is determined using a Mann-Whitney test. **(B)** Heatmap showing pair-wise comparison of spatial association scores between sub-clinical and clinical groups. The proximity of each cell population to other populations indicated in B3-type lesions were determined using 50 μm radius range.

**Table S1.** Clinical information.

| ID | Sex | Age | AFB<sup>a</sup> | GeneXpert | Sputum Culture | GMS<sup>b</sup> | PAS<sup>c</sup> | HRZE | TB Treatment Duration | MDR DST<sup>d</sup> | Time since recording | Pathology Presentation | Location |
| --- | --- | --- | --- | --- | --- | --- | --- | --- | --- | --- | --- | --- | --- |
| 1 | M | 28 | P<sup>e</sup> | N<sup>f</sup> | N | N | N | + | 2mo | N | 5mo | Caseous Necrosis | Left upper lobe |
| 2 | M | 58 | P | P | N | N | N | - | N/A | N | 1d | Caseous Necrosis | Left upper lobe |
| 3 | F | 34 | P | P | ND<sup>g</sup> | N | N | + | 8d | N | 1mo | Caseous Necrosis | Left lower lobe |
| 4 | M | 33 | N | P | N | N | N | After<sup>h</sup> | 1mo | N | 2yr | Caseous Necrosis | Right upper lobe |
| 5 | F | 20 | N | ND | P | N | N | + | 5mo | ND | 9mo | Caseous Necrosis | Right upper lobe |
| 6 | M | 33 | P | ND | ND | N | ND | + | 1mo | ND | 1mo | Tuberculoma | Left upper lobe |
| 7 | M | 33 | P | P | N | N | N | + | 4mo | N | 1mo | Caseous Necrosis | Left lower lobe |
| 8 | F | 29 | N | ND | N | N | ND | + | 20mo | ND | 5yr | Caseous Necrosis | Right upper lobe |
| 9 | F | 48 | N | P | P | N | N | + | 9mo | ND | 15mo | Caseous Necrosis | Right mid lower lobe |
| 10 | M | 48 | N | P | N | N | N | + | 1mo | ND | 1mo | Coagulative necrosis, MGCs | Left upper lobe |
| 11 | F | 25 | N | ND | P | N | ND | + | 12mo | P | 2yr | Tuberculosis-destructed lungs | Right mid lower lobe |
| 12 | M | 44 | ND | ND | ND | ND | ND | After | ND | ND | 6mo | Tuberculoma | Left upper lobe |
| 13 | M | 39 | ND | ND | ND | ND | ND | After | ND | ND | 1yr | Tuberculosis | Right upper lobe |
| 14 | M | 56 | ND | ND | ND | ND | ND | After | ND | ND | 2yr | Caseous necrosis, MGCs | Left lower lobe |
| 15 | M | 56 | Pt<sup>i</sup> | ND | ND | ND | ND | After | ND | ND | 1yr | Caseous necrosis, MGCs | Right upper lobe |
| 16 | M | 53 | ND | ND | ND | ND | ND | After | ND | ND | 1yr | Tuberculosis | Right upper lobe |
| 17 | M | 61 | ND | ND | ND | ND | ND | After | ND | ND | 6mo | granulomatous inflammation | Left lower + Right upper lobe |
| 18 | M | 42 | Pt | ND | ND | N | N | After | ND | ND | 1yr | Caseous necrosis | Right upper lobe |
| 19 | M | 40 | ND | ND | ND | ND | ND | After | ND | ND | 2yr | Tuberculoma | Right lower lobe |
| 20 | M | 45 | N | ND | ND | N | N | After | ND | ND | 3yr | Caseous necrosis, MGCs | Left upper lobe |
| 21 | M | 54 | Pt | ND | ND | ND | ND | After | ND | ND | 1yr | Coagulative necrosis, MGCs | Left upper lobe |
| 22 | F | 48 | ND | ND | ND | ND | ND | After | ND | ND | 6yr | granulomatous inflammation | Left upper lobe (partial) |

a. Acid-fast bacteria
b. Grocott’s methenamine silver
c. Periodic acid-Schiff
d. Multidrug-resistant drug sensitivity test
e. Positive
f. Negative
g. Not Determined
h. Therapy after resection
i. Positive AFB in tissue

**Table S2.**
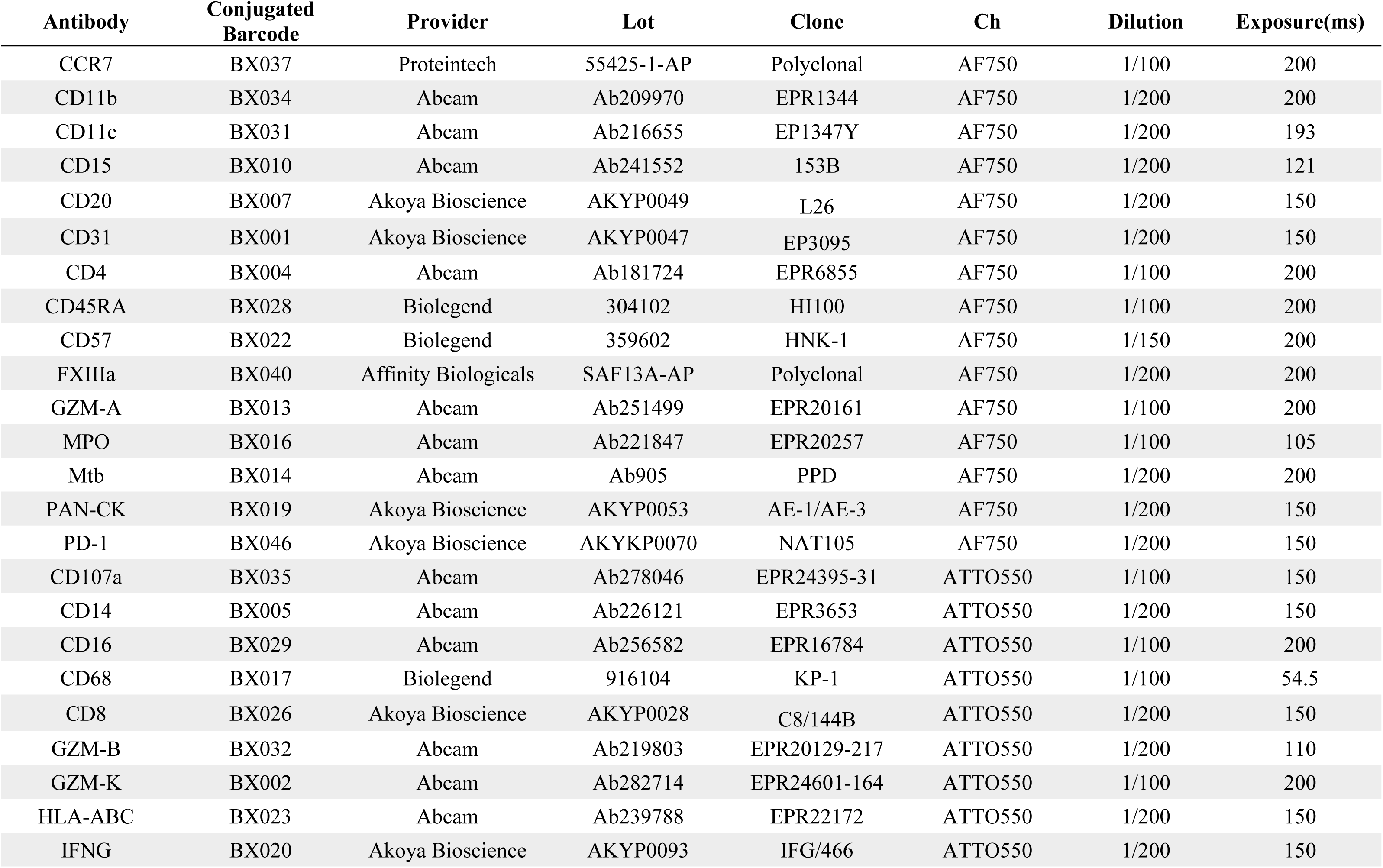
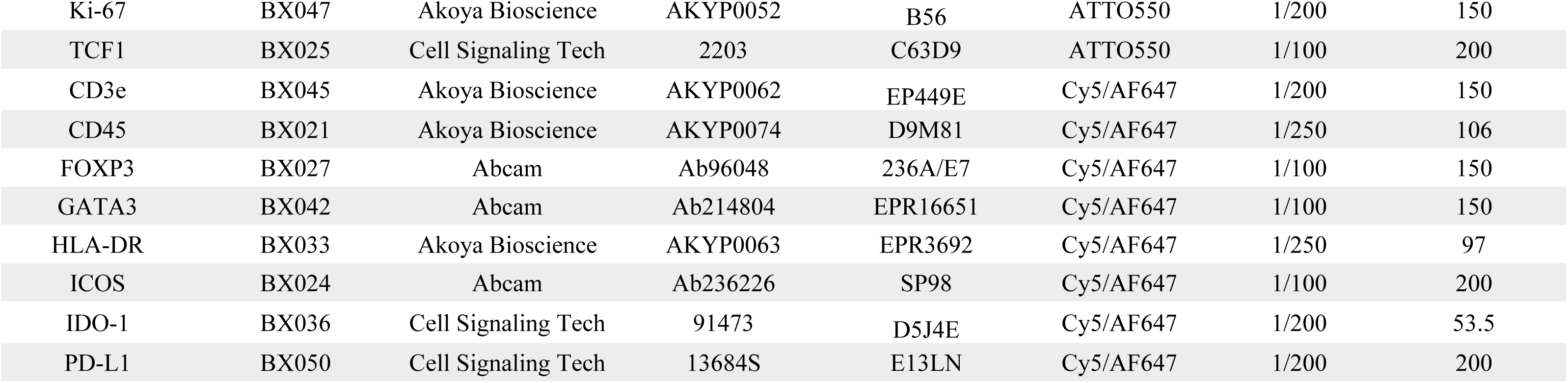
Antibody list and condition for PCF staining.

| **Antibody** | **Conjugated Barcode** | **Provider** | **Lot** | **Clone** | **Ch** | **Dilution** | **Exposure(ms)** |
| --- | --- | --- | --- | --- | --- | --- | --- |
| CCR7 | BX037 | Proteintech | 55425-1-AP | Polyclonal | AF750 | 1/100 | 200 |
| CD11b | BX034 | Abcam | Ab209970 | EPR1344 | AF750 | 1/200 | 200 |
| CD11c | BX031 | Abcam | Ab216655 | EP1347Y | AF750 | 1/200 | 193 |
| CD15 | BX010 | Abcam | Ab241552 | 153B | AF750 | 1/200 | 121 |
| CD20 | BX007 | Akoya Bioscience | AKYP0049 | L26 | AF750 | 1/200 | 150 |
| CD31 | BX001 | Akoya Bioscience | AKYP0047 | EP3095 | AF750 | 1/200 | 150 |
| CD4 | BX004 | Abcam | Ab181724 | EPR6855 | AF750 | 1/100 | 200 |
| CD45RA | BX028 | Biolegend | 304102 | HI100 | AF750 | 1/100 | 200 |
| CD57 | BX022 | Biolegend | 359602 | HNK-1 | AF750 | 1/150 | 200 |
| FXIIIa | BX040 | Affinity Biologicals | SAF13A-AP | Polyclonal | AF750 | 1/200 | 200 |
| GZM-A | BX013 | Abcam | Ab251499 | EPR20161 | AF750 | 1/100 | 200 |
| MPO | BX016 | Abcam | Ab221847 | EPR20257 | AF750 | 1/100 | 105 |
| Mtb | BX014 | Abcam | Ab905 | PPD | AF750 | 1/200 | 200 |
| PAN-CK | BX019 | Akoya Bioscience | AKYP0053 | AE-1/AE-3 | AF750 | 1/200 | 150 |
| PD-1 | BX046 | Akoya Bioscience | AKYKP0070 | NAT105 | AF750 | 1/200 | 150 |
| CD107a | BX035 | Abcam | Ab278046 | EPR24395-31 | ATTO550 | 1/100 | 150 |
| CD14 | BX005 | Abcam | Ab226121 | EPR3653 | ATTO550 | 1/200 | 150 |
| CD16 | BX029 | Abcam | Ab256582 | EPR16784 | ATTO550 | 1/100 | 200 |
| CD68 | BX017 | Biolegend | 916104 | KP-1 | ATTO550 | 1/100 | 54.5 |
| CD8 | BX026 | Akoya Bioscience | AKYP0028 | C8/144B | ATTO550 | 1/200 | 150 |
| GZM-B | BX032 | Abcam | Ab219803 | EPR20129-217 | ATTO550 | 1/200 | 110 |
| GZM-K | BX002 | Abcam | Ab282714 | EPR24601-164 | ATTO550 | 1/100 | 200 |
| HLA-ABC | BX023 | Abcam | Ab239788 | EPR22172 | ATTO550 | 1/200 | 150 |
| IFNG | BX020 | Akoya Bioscience | AKYP0093 | IFG/466 | ATTO550 | 1/200 | 150 |
| Ki-67 | BX047 | Akoya Bioscience | AKYP0052 | B56 | ATTO550 | 1/200 | 150 |
| TCF1 | BX025 | Cell Signaling Tech | 2203 | C63D9 | ATTO550 | 1/100 | 200 |
| CD3e | BX045 | Akoya Bioscience | AKYP0062 | EP449E | Cy5/AF647 | 1/200 | 150 |
| CD45 | BX021 | Akoya Bioscience | AKYP0074 | D9M81 | Cy5/AF647 | 1/250 | 106 |
| FOXP3 | BX027 | Abcam | Ab96048 | 236A/E7 | Cy5/AF647 | 1/100 | 150 |
| GATA3 | BX042 | Abcam | Ab214804 | EPR16651 | Cy5/AF647 | 1/100 | 150 |
| HLA-DR | BX033 | Akoya Bioscience | AKYP0063 | EPR3692 | Cy5/AF647 | 1/250 | 97 |
| ICOS | BX024 | Abcam | Ab236226 | SP98 | Cy5/AF647 | 1/100 | 200 |
| IDO-1 | BX036 | Cell Signaling Tech | 91473 | D5J4E | Cy5/AF647 | 1/200 | 53.5 |
| PD-L1 | BX050 | Cell Signaling Tech | 13684S | E13LN | Cy5/AF647 | 1/200 | 200 |

## Notes

### Competing Interest Statement

The authors have declared no competing interest.

